# Polar chromosomes are rescued from missegregation by spindle elongation-driven microtubule pivoting

**DOI:** 10.1101/2024.06.16.599200

**Authors:** Isabella Koprivec, Valentina Štimac, Mario Đura, Kruno Vukušić, Petra Mikec, Iva M. Tolić

**Author notes:** These authors contributed equally to this work (Isabella Koprivec, Valentina Štimac).

## Abstract

Polar chromosomes, which initially attach to the mitotic spindle behind the pole, are prone to missegregation and micronuclear entrapment, contributing to chromosomal instability in cancer. Yet, the mechanisms ensuring their faithful segregation remain unclear. Here, we show that polar chromosomes require a unique step involving spindle elongation, which repositions chromosome-bound astral microtubules by pivoting them around the centrosome toward the spindle surface. By modulating Eg5/KIF11 activity, we demonstrate that spindle elongation determines the direction and extent of pivoting, with microtubules from the opposite spindle half facilitating final movement. Kinetochores on polar chromosomes form mixed lateral and immature monotelic attachments, recruiting corona components and partially Mad2, but lacking Astrin. In several cancer cell lines, limited spindle elongation delays polar chromosome resolution, whereas enhanced elongation accelerates it. These findings highlight the role of spindle elongation in the timely rescue of polar chromosomes from the ‘danger zone’ behind the pole, and provide mechanistic insight into how chromosome congression errors can arise in cancer.

## Introduction

Chromosomes must efficiently congress to the equatorial plane of the spindle to ensure proper segregation (Maiato et al., 2017). Based on their position at the nuclear envelope breakdown (NEBD), chromosomes undertake different pathways toward the equator, resulting in distinct congression efficiencies and segregation error rates (Vukušić and Tolić, 2022). Chromosomes behind the spindle poles are particularly at risk of missegregation, as they are shielded by the pole and require a multi-step process to reach the equator (Barisic et al., 2014; Klaasen et al., 2022). First, these chromosomes approach the spindle pole from the back through centripetal movement driven by kinetochore dynein pulling along astral microtubules and nuclear actomyosin (Barisic et al., 2014; Booth et al., 2019; Li et al., 2007; Rieder and Alexander, 1990; Yang et al., 2007). Subsequently, the chromosomes travel along the main spindle surface using CENP-E at kinetochores, pulling forces by end-on attached microtubules, and polar ejection forces (Barisic et al., 2014; Cai et al., 2009; Craske and Welburn, 2020; Kapoor et al., 2006; McEwen et al., 2001; Renda et al., 2022; Shrestha and Draviam, 2013; Vukušić and Tolić, 2024; Wandke et al., 2012). However, the mechanism by which polar chromosomes traverse the pole and shift from the astral region to the main spindle remains unknown.

Chromosomes that fail to escape the area behind the pole before the spindle fully forms risk stabilization of improper attachments, ensheathment by the endoplasmic reticulum, and missegregation in case of the spindle assembly checkpoint (SAC) weakness, bypass or delayed satisfaction (Dick and Gerlich, 2013; Ferrandiz et al., 2022; Gomes et al., 2022; Kabeche and Compton, 2013; Klaasen et al., 2022). Recent studies discovered that such unaligned (i.e., behind the spindle pole) and misaligned (i.e., near the pole on the main spindle) chromosomes represent an important source of segregation errors, with the highest probability of ending up in micronuclei, the strongest correlation with chromosomal instability, and a direct link to metastatic potential (Gomes et al., 2022; Tucker et al., 2023). Several labs have observed mis- and unaligned chromosomes in cancer cell lines, such as ovarian, bone, colorectal and breast cancer, and in patient samples of primary and metastatic breast cancer (Ertych et al., 2014; Gomes et al., 2022; Tamura et al., 2020; Tucker et al., 2023). Multiple lines of evidence also indirectly suggest that chromosome 1, whose 1q arm gain directly leads to cancer development and malignant transformation, might often be located behind the pole and thus at risk of unalignment (Bolzer et al., 2005; Girish et al., 2023; Klaasen et al., 2022; Tovini and McClelland, 2019). While these implications highlight the importance of escaping the polar region on time, the reason why chromosomes occasionally fail to do so, especially in cancer cells, is still unclear.

Here, we develop approaches integrating stimulated emission depletion (STED), lattice light-sheet (LLS), and confocal microscopy with molecular perturbations to overcome spatiotemporal resolution barriers that hinder studies of polar chromosome dynamics, which occurs quickly within a spindle region that is densely populated with microtubules. We present a mechanism crucial for chromosome segregation, based on a fundamental, yet previously unexplored, property of centrosome-anchored microtubules—their ability to pivot around the centrosome. Our study reveals that spindle elongation prompts microtubules attached to polar chromosomes to pivot towards the center of the spindle, releasing the chromosomes from the "danger zone" behind the spindle pole and enabling their contact with the spindle surface. During pivoting, polar chromosomes form predominantly mono-lateral and other complex attachments with astral microtubules, which persist until the chromosome contacts the spindle surface and contribute to spindle assembly. From a functional perspective, we demonstrate a critical role of spindle elongation-driven pivoting in promoting timely mitosis and preventing chromosome alignment errors in cancer cells, suggesting potential strategies to modulate these errors.

## Results

### Polar chromosomes require a unique congression step to transit across the spindle pole

Given that chromosomes located behind the spindle pole at NEBD take longer to congress to the metaphase plate in comparison to other chromosomes (Klaasen et al., 2022), we hypothesized that they require a specific congression step. Alternatively, they may exhibit slower progression of the steps shared with other chromosomes. To distinguish between these possibilities, we imaged human retinal pigment epithelium (RPE1) cells with labeled centromeres and either tubulin or centrosomes using a synergy of STED immunofluorescence (Figure 1A) and LLS live-cell imaging (Figure 1B). STED microscopy enabled visualization of microtubules within the polar region in super-resolution (50-100 nm, Supplementary Figure 1A) (Koprivec et al., 2025). This level of resolution was essential for distinguishing individual microtubules or small bundles together with their attachments to kinetochores within a microtubule-packed region, considering that human spindles contain about 6000 microtubules (Kiewisz et al., 2022). LLS microscopy enabled imaging of all kinetochores throughout mitosis at a time resolution of 6 seconds due to the low phototoxicity of this technique (Shi et al., 2024). The high time resolution was crucial for tracking individual polar kinetochores, as they move quickly and pass closely by other kinetochores (Video 1).

**Figure 1.**
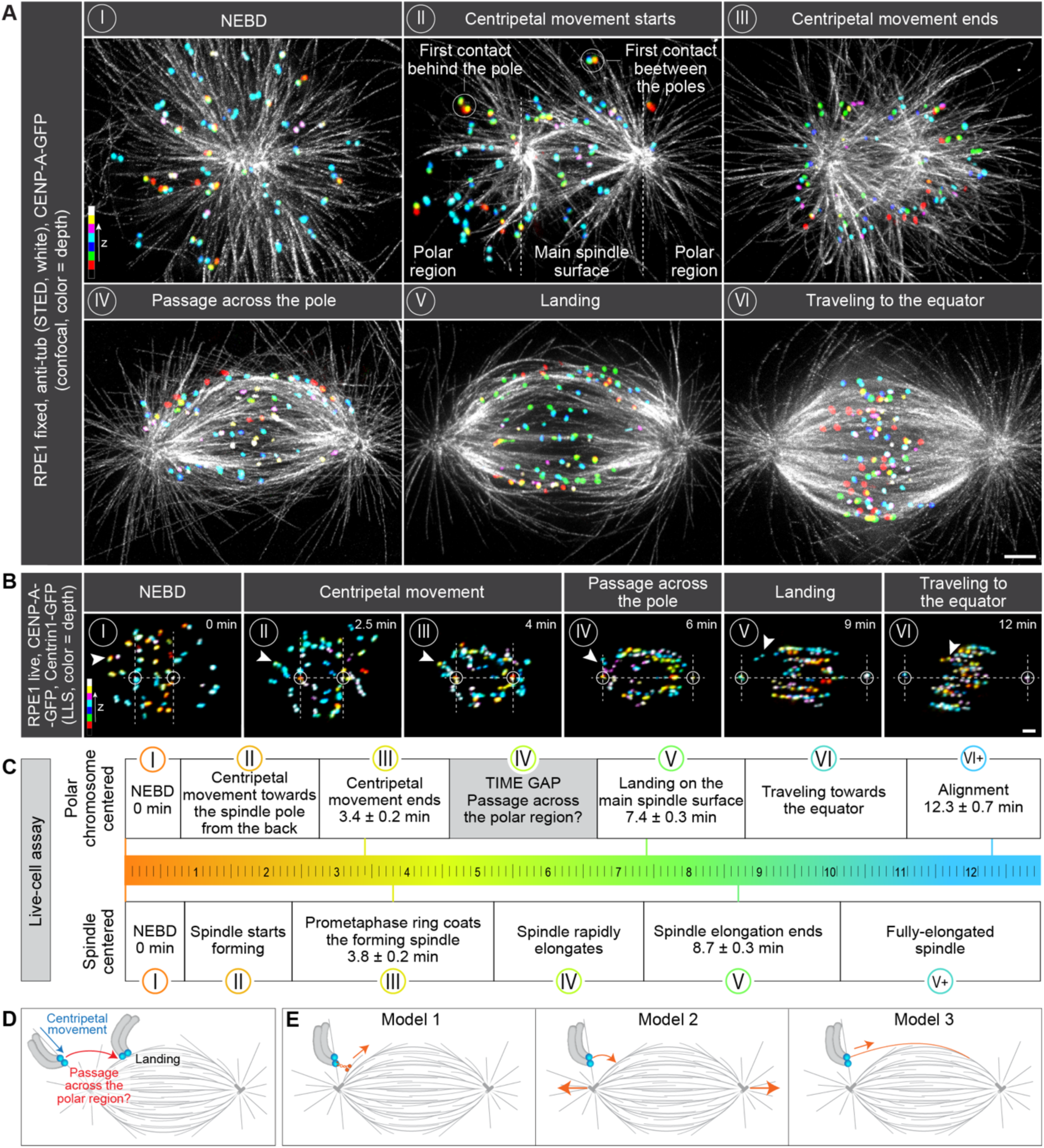
Polar chromosomes require an additional congression step to pass the polar region. **(A)** STED superresolution images of microtubules immunostained for α-tubulin (white) in RPE1 cells stably expressing CENP-A-GFP (colorful, confocal). Images show maximum intensity projections of the whole cell. Kinetochores are color-coded for depth with the brgbcmyw LUT in ImageJ throughout 8-12 z-planes in prometaphase and 35 z-planes in metaphase, corresponding to 2.4-3.6 µm and 10.5 µm, respectively. The images represent different phases of mitosis: I – nuclear envelope breakdown (NEBD), II – a spindle starts forming and kinetochores in the polar region start their centripetal movement towards the spindle pole from the back, III – a prometaphase ring forms and centripetal movement of kinetochores ends, IV – a spindle rapidly elongates while kinetochores located at the back of the pole pass the polar region, V – spindle elongation ends and all kinetochores land onto the spindle surface, VI – a spindle is fully elongated and kinetochores travel towards the spindle equator. These stages were inferred from the arrangements of microtubules and kinetochores, see Methods. The polar region and the main spindle surface are denoted in image II. **(B)** Time-lapse images of RPE1 cells stably expressing CENP-A-GFP and Centrin1-GFP (colorful, lattice light-sheet). Images show maximum intensity projections of the whole cell. Kinetochores are color-coded for depth with the brgbcmyw LUT in ImageJ throughout 32-40 z-planes, corresponding to 4.6-5.8 µm. Mitotic phases from I to VI correspond to the ones described in (A). The polar region is denoted as a region behind two dashed lines perpendicular to the long spindle axes. Centrosomes are denoted with white circles. White arrows represent one polar kinetochore pair tracked from NEBD until alignment to the spindle equator. **(C)** Polar chromosome-centered (upper) and spindle-centered (lower) timelines of prometaphase events. Events are marked with roman numerals that correspond to the ones shown in (A) and (B). The times are calculated based on 39 kinetochore pairs in 20 cells from 8 experiments. **(D)** Schematic representation of the first (end of centripetal movement) and the last (landing) known time points between which the time gap and possible passage across the polar region occur. **(E)** Three possible models for passing the spindle pole. In the first model, a motor bound to polar kinetochores brings them onto the spindle surface, representing diverse mechanisms based on kinetochore- and chromosome-bound motors. The second model proposes that spindle elongation drives pivoting of astral microtubules that consequently bring polar kinetochores in front of the spindle pole. In the last model, polar kinetochores attach to microtubules emanating from the opposite spindle half that bring them onto the spindle surface. All scale bars, 2 µm.

By using LLS movies (Figure 1B), we found that at NEBD, 15 ± 2 chromosomes per cell (all data are given as mean ± SEM, n = 24 cells) were located behind the spindle poles (Supplementary Figure 1B-D). Out of those 15, 8 ± 1 ended their centripetal movement in the region between the two poles, whereas 7 ± 1 chromosomes ended their centripetal movement in the region behind the pole (see examples in Figure 1A and Supplementary Figure 1B). We refer to the latter group as polar chromosomes. By constructing a timeline of the polar chromosome movements (Figure 1B-C, see individual stages inferred from the arrangements of microtubules and kinetochores in Figure 1A), we found that after NEBD a typical polar chromosome started its centripetal movement approaching the spindle pole from the back until it reached a relatively stable distance from the pole at 3.4 ± 0.2 minutes, which roughly corresponds to the formation of the prometaphase chromosome ring around the nascent spindle (Magidson et al., 2011). Remarkably, it took additional 4 minutes for polar chromosomes to reach the spindle surface, an event we term "landing", which occurred 7.4 ± 0.3 minutes after NEBD (Figure 1B-C). During this time gap, the spindle rapidly elongated and other chromosomes have already started congressing to the metaphase plate. Once the polar chromosomes landed on the spindle surface, they immediately started traveling towards the spindle midplane, finishing aligning at 12.3 ± 0.3 minutes (Figure 1B-C, Supplementary Figure 1E).

The observed time gap may be specific to the location of polar chromosomes behind the pole in prometaphase, but it may also be due to their large distance to the spindle midplane. To discriminate between these possibilities, we compared the timing of polar and non-polar chromosomes with a similar initial distance to the centrosome midpoint (Supplementary Figure 1F), and found that only polar chromosomes showed a delay in landing and a time gap from the end of the centripetal movement to the attachment to the spindle surface, indicating that the gap is specific to chromosome location (Supplementary Figure 1G-H). The other segments of the chromosome’s travel, the centripetal movement and the transfer along the spindle surface, lasted similarly for polar and non-polar chromosomes (Supplementary Figure 1I-J), which argues against polar chromosomes having slower progression of shared steps. Thus, the delayed alignment of polar chromosomes causing their susceptibility to missegregation is due to a time gap wherein the polar chromosome traverses across the pole to reach the main spindle surface. This reveals an additional step in congression specific to polar chromosomes (Figure 1D).

To test whether this additional step includes the attachments to microtubules, we analyzed STED images (Figure 1A) and found that 58.1 ± 3.2 % of chromosomes already had microtubules next to their kinetochores at a time close to NEBD, whereas 41.9 ± 3.2 % were completely unattached (Supplementary Figure 1K-L). By the end of the centripetal movement phase all kinetochore pairs were attached to microtubules and remained attached throughout the subsequent phases (Supplementary Figure 1K-L). Some of the chromosomes that were unattached and located behind the pole at NEBD might have remained unattached as they were passed by the moving centrosome, resulting in their first contact with spindle microtubules between the spindle poles. In contrast, all polar chromosomes, which made the first contact behind the pole, were attached to microtubules during the transit across the pole. Thus, pole crossing by the polar chromosomes involves the attachment of kinetochores to microtubules.

### Passage across the pole occurs independently of the motor proteins implicated in chromosome congression

To uncover the mechanism of the passage across the polar region, we envisioned three models (Figure 1E). In the first model, motor proteins located at the kinetochore (CENP-E or dynein) or on the chromosome arm (chromokinesins) of polar chromosomes attach to the microtubules of the main spindle surface and pull the polar chromosome in the direction of the spindle surface. In the second model, the spindle elongates, causing both poles to move outward, thereby repositioning chromosomes from the back of the spindle pole to the main spindle surface. In the third model, microtubules extending from the opposite side of the spindle attach to the polar chromosome and pull it towards the main spindle surface.

To test the first model, we depleted CENP-E (kinesin-7) (Barisic et al., 2014; Cai et al., 2009; Kapoor et al., 2006; Maiato et al., 2017), Spindly, which promotes dynein recruitment to kinetochores in early mitosis (Griffis et al., 2007), or co-depleted the chromokinesins KIF4A (kinesin-4) and Kid (kinesin-10) (Mazumdar et al., 2004; Wandke et al., 2012; Yajima et al., 2003) by using small interfering RNAs (siRNAs) (Supplementary Figure 2A). As an alternative approach to disrupt CENP-E activity, we used the small-molecule inhibitor GSK923295 (Wood et al., 2010) (Supplementary Figure 2A). We also perturbed actomyosin contraction (Booth et al., 2019) by depolymerizing actin via Latrunculin A (Supplementary Figure 2A), thereby completing the test of motor-driven mechanisms that were previously implied in chromosome congression (Maiato et al., 2017). Even when those proteins were effectively depleted or inhibited, as determined by a ∼90 % reduction of their signal, absence from the expected location, and the characteristic phenotype (Supplementary Figure 2B-C), nearly all polar chromosomes successfully traveled from the back of the spindle towards the main spindle surface (Supplementary Figure 2D). A tiny fraction (∼2 %) of polar chromosomes after Spindly depletion remained behind the pole, as they were ejected more than 6 µm away from the pole into the region beyond the future area of the spindle (Supplementary Figure 2D-E). In contrast, in control cells, polar kinetochores approached the pole and remained at a distance of about 3 µm, indicating the importance of proximity to the pole for successful passage across the pole (Supplementary Figure 2E). Given that Spindly and CENP-E are essential for the expansion of the kinetochore fibrous corona (Pereira et al., 2018; Sacristan et al., 2018; Wu et al., 2023), these data show that the expanded corona is dispensable for the initial chromosome transit across the pole. Taken together, these results show that the passage across the pole occurs independently of the previously described congression drivers, but benefits from the role of dynein in keeping the chromosomes close to the spindle pole.

### Pivoting of astral microtubules around the centrosome repositions polar chromosomes close to the spindle surface

In the second model of polar chromosome resolution (Figure 1E), spindle elongation leads to repositioning of polar chromosomes attached to astral microtubules towards the main spindle surface. This model relies on two assumptions: 1) chromosome-attached microtubules are able to pivot around the centrosome, 2) chromosomes move through the cytoplasm considerably less than the centrosomes. To explore these assumptions, we tracked in 3D the polar kinetochores and their microtubules, as well as centrosomes and chromosome arms, from the end of the centripetal movement until after landing on the spindle surface (Figure 2A-C, Supplementary Figure 2F, Video 2-4). To quantify the pivoting motion of a polar chromosome, we measured the angle between the centrosome-centrosome axis and the line connecting the sister kinetochores midpoint with the centrosome, termed "pivoting angle" (Figure 2D). The pivoting angle showed a fast decrease during the period of rapid spindle elongation and stabilized as the spindle length stabilized (Figure 2E-F). Moreover, the pivoting angle was strongly anti-correlated with the spindle length for individual kinetochore pairs (Figure 2G, Supplementary Figure 2G). These results indicate that polar kinetochores, along with their attached microtubules, pivot around the centrosome towards the spindle center.

**Figure 2.**
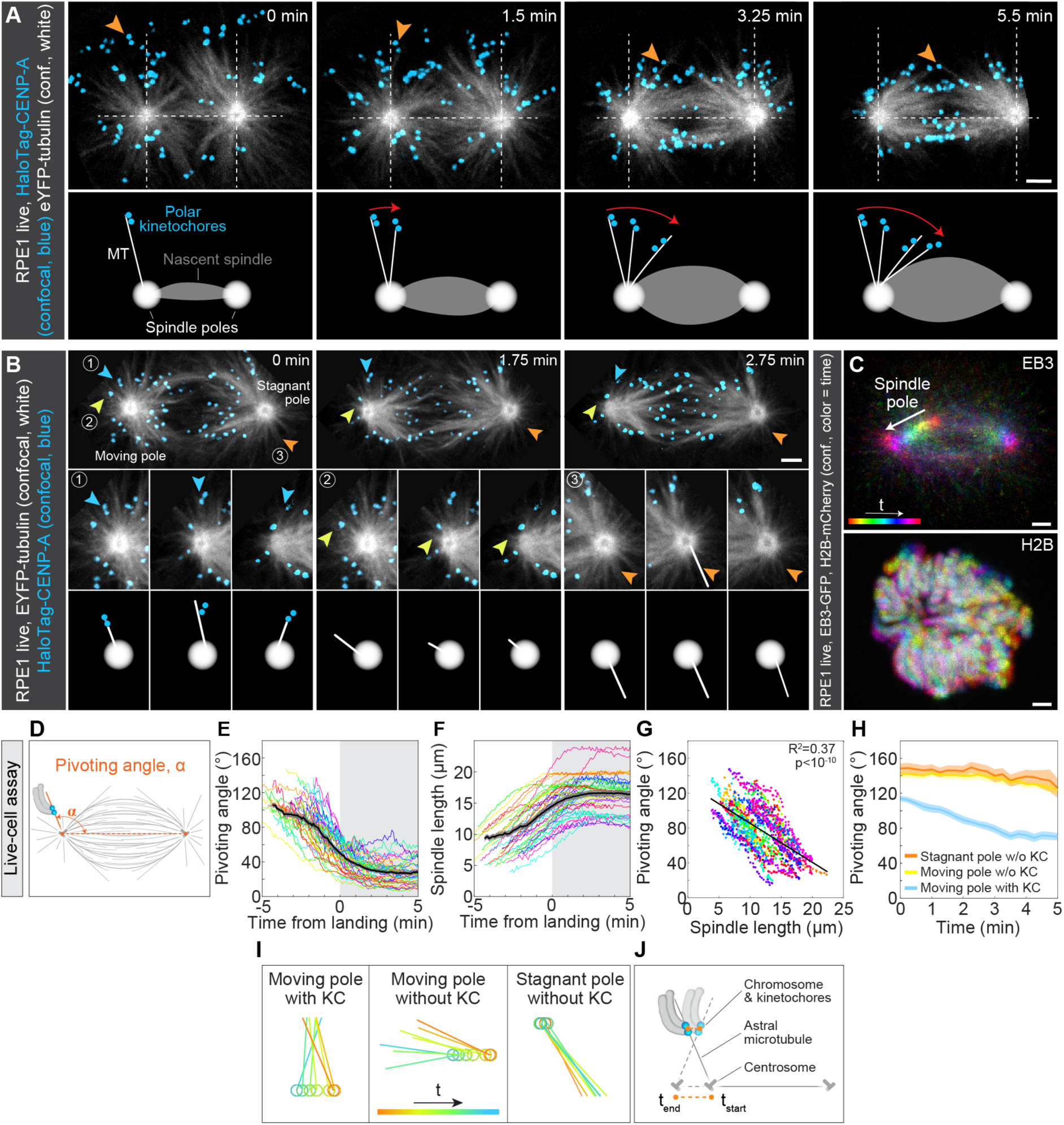
Polar chromosomes rely on spindle elongation-driven pivoting of astral microtubules to pass the polar region. **(A)** Upper panel: time-lapse images of RPE1 cells stably expressing EYFP-tubulin (white, confocal) and H2B-mRFP (not imaged) with added HaloTag-CENP-A (blue, confocal). Each image represents a single central z-plane and was rotated and cropped to align the spindle horizontally and position it in the center of the frame. The orange arrow denotes the kinetochore pair attached to the pivoting astral microtubule. Lower panel: Schematic representation of pivoting (red arrow) based on the images from the upper panel, showing the current and previous microtubule positions. **(B)** Time-lapse images of RPE1 cells stably expressing EYFP-tubulin (white, confocal) and H2B-mRFP (not imaged) with added HaloTag-CENP-A (blue, confocal). Moving and stagnant spindle poles are denoted in the first image. The blue arrow represents the microtubule attached to the polar kinetochore tracked over time. Yellow and orange arrows represent microtubules without attached polar kinetochores on the moving and stagnant spindle pole, respectively. Insets and schematic representations of the previously described microtubule types are shown below the images of the whole spindle; spindle poles are depicted as white circles. Images are maximum intensity projections of 3 central z-planes, corresponding to 3 µm. **(C)** Images of RPE1 cells stably expressing EB3-GFP (colorful, confocal, upper image) and H2B-mCherry (colorful, confocal, lower image). Images are temporal maximum intensity projections of the centrosome (upper) and chromosome (lower) movements over 3 central z-planes (3 µm) and a time period of 3.3 minutes, color-coded for time with the Spectrum LUT in ImageJ. Spindle pole movement is depicted with white arrow. **(D)** Schematic representation of measured parameters for polar kinetochores along with the starting (end of centripetal movement) and last (landing) tracking point (t = 0 minutes). Spindle length was determined as the distance between the two spindle poles. The pivoting angle was determined as an angle that the midpoint of the kinetochore pair closes with the long spindle axis. **(E)** The pivoting angle of polar kinetochore pairs and **(F)** corresponding spindle length in time. The landing point of each kinetochore pair is set to zero. Values are shown as mean (dark line) and SEM (shaded areas). N = 39 kinetochore pairs in 20 cells from 8 experiments. **(G)** Anticorrelation of the pivoting angle and spindle length for the same polar kinetochores as in (E). Each color represents one polar kinetochore pair and each dot represents its pivoting angle and the corresponding spindle length in time. Linear regression. N = 39 kinetochore pairs in 20 cells from 8 experiments. **(H)** The pivoting angle of the microtubule attached to the kinetochores on the moving pole (blue), microtubule without attached kinetochores on the moving pole (yellow) and microtubule without attached kinetochores on the stagnant pole (orange) in time. Values are shown as mean (dark line) and SEM (shaded areas). N = 16 microtubules with polar kinetochores attached to them on the moving pole in 10 cells from 4 experiments. N = 16 microtubules without polar kinetochores attached to them on the moving pole in 10 cells from 4 experiments. N = 14 microtubules without polar kinetochores attached to them on the stagnant pole in 6 cells from 4 experiments. The fourth group, microtubules with polar kinetochores attached to them on the stagnant pole, could not be found, as poles were stagnant only when they were already at the nuclear border at NEBD and, therefore, had no polar chromosomes to begin with. **(I)** Schematic representation of the spindle pole movement and pivoting angle for each microtubule marked in (B) in time (from red to blue). Note that only the microtubule attached to polar kinetochores pivots around the spindle pole. **(J)** Schematic representation of the centrosome, astral microtubule and chromosome movements during pivoting. Note that the centrosome moves significantly while microtubules pivot under the weight of a chromosome which moves only slightly. All scale bars, 2 µm.

To directly see the pivoting motion of astral microtubules, which are undetectable in typical videos of mitosis, we developed a confocal microscopy protocol with adjusted pixel size (See Methods) to image RPE1 cells expressing EYFP-α-tubulin with added HaloTag-CENP-A (Figure 2A, Video 2 and 3). The kinetochores of a polar chromosome were found to be adjacent to a microtubule signal representing a single astral microtubule, or a few in a bundle. We found that this astral microtubule pivoted around the centrosome in the direction of the spindle surface, while maintaining its connection to the polar chromosome, during the centrosome’s movement outwards (Figure 2A and 2H-I). The microtubules remained straight instead of bending (Supplementary Figure 2H), indicating that the entire microtubule pivots around the centrosome, as opposed to solely the plus-end-proximal segment gliding along the cell cortex or organelles. Overall, these direct observations of microtubule pivoting support the second model and indicate that the force driving the pivoting of chromosome-attached microtubules exceeds the force preserving their extension direction at the centrosome.

In contrast to the chromosome-carrying astral microtubules, the microtubules that were not attached to a chromosome did not exhibit directed pivoting but instead pivoted around the centrosome in a random manner (Figure 2B and 2H-I, Supplementary Figure 2I-J). Their pivoting was similar in the cases when their centrosome moved significantly and when it appeared stationary (Figure 2B and 2H-I, Supplementary Figure 2I-K). Stationary poles were observed in cells where one pole was positioned at the nuclear border during NEBD, resulting in the absence of polar chromosomes and the presence of only chromosome-free microtubules at this pole. Whereas chromosome-carrying microtubules pivoted by ∼36°, on average, in 3 minutes, the chromosome-free microtubules changed their angle by less than ∼8° irrespective of the pole movement (Supplementary Figure 2I). These findings suggest that the force driving the pivoting of chromosome-free microtubules is independent of the centrosome movement and smaller than the force maintaining their direction at the centrosome.

For spindle elongation to reposition polar chromosomes towards the main spindle surface, chromosomes must move less than centrosomes during this process. This was confirmed by temporal maximum-intensity projections of spindles labeled with EB3-GFP and H2B-mCherry, which showed that chromosomes remained relatively stationary while the centrosomes moved outward (Figure 2C, Video 4). Collectively, these findings support the model in which spindle elongation induces pivoting of astral microtubules attached to stationary polar chromosomes (Figure 2J), enabling their interaction with the spindle surface.

### Direction of the centrosome movement dictates the direction of microtubule pivoting

The pivoting model provides a straightforward prediction: if the spindle dynamics reverses from elongation to shortening, the direction of microtubule pivoting should also reverse, implying that microtubules should pivot towards the region behind the spindle pole instead of towards the main spindle surface. Similarly, a constant spindle length should be associated with negligible pivoting. To test these predictions, we designed a set of manipulations of Eg5/KIF11 (kinesin-5), the main motor protein driving spindle elongation (Kapitein et al., 2005; Kashina et al., 1997), to either reverse, restore, or block spindle elongation (Figure 3A-D, Video 5 and 6).

**Figure 3.**
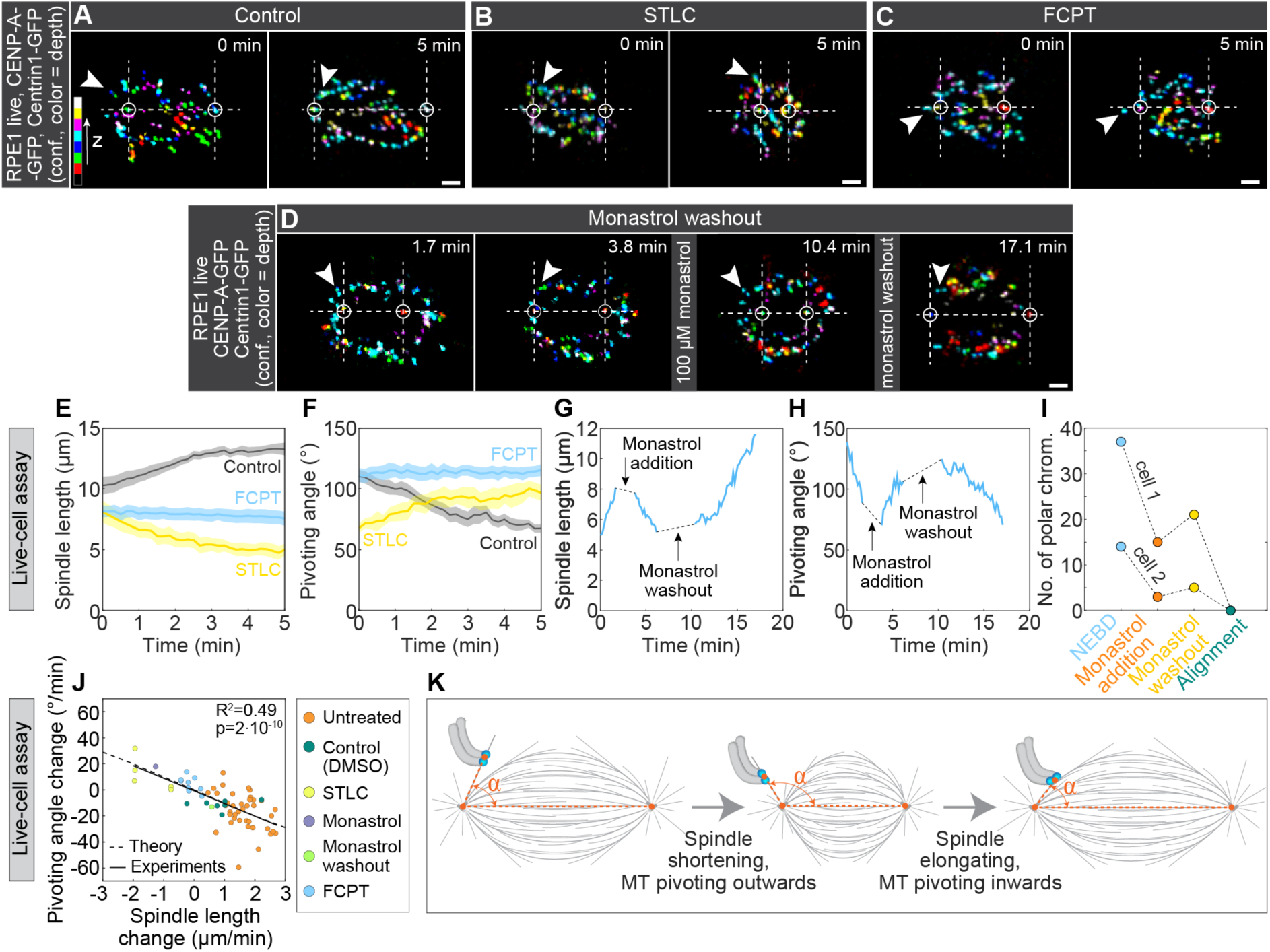
Centrosome movement determines the direction and magnitude of pivoting. **(A-D)** Time-lapse images of RPE1 cells stably expressing CENP-A-GFP and Centrin1-GFP (colorful, confocal) for (A) DMSO control, (B) STLC-, (C) FCPT-, and (D) monastrol-treated cells. White arrows represent one polar kinetochore pair over time. White circles represent spindle poles, horizontal dashed lines represent the long spindle axis and lines perpendicular to it denote the borders of the polar region. Note that the STLC treatment caused spindle shortening, FCPT treatment blocked spindle elongation, whereas monastrol treatment caused spindle shortening that could be reversed by washout. Kinetochores are color-coded for depth with the brgbcmyw LUT in ImageJ throughout 14-21 z-planes, corresponding to 5.6-8.4 µm. **(E)** Spindle length and **(F)** pivoting angle in time for polar kinetochores in control (gray), STLC (yellow) and FCPT (blue) treatment. Values are shown as mean (dark line) and SEM (shaded areas). t = 0 minutes corresponds to inhibitor addition and activity. N = 7 kinetochore pairs in 7 cells from 7 experiments (DMSO), N = 5 kinetochore pairs in 3 cells from 3 experiments (STLC), N = 9 kinetochore pairs in 7 cells from 7 experiments (FCPT). **(G)** Spindle length and **(H)** pivoting angle in time during the monastrol washout experiment, for the spindle and kinetochore pair marked in (D). Dashed lines denote the time required for the addition or washout of monastrol together with the time required for the spindle length to start changing. **(I)** Number of chromosomes behind the pole at the indicated times in the monastrol washout experiment for Cell 1 (shown in (D)) and Cell 2 (shown in Supplementary Figure 3D). **(J)** The experimental correlation of the pivoting angle change and spindle length change for the control (orange), DMSO control (dark green), monastrol washout (light green), FCPT (blue), monastrol (purple) and STLC (yellow) treatments along with the theoretical curve calculated based on the experimental parameters from DMSO and STLC experiments (See Supplementary Figure 3). Linear regression. **(K)** Schematic representation of pivoting angle change based on the spindle length change. All scale bars, 2 µm.

To reverse spindle elongation into shortening, we added the Eg5 inhibitor STLC (S-Trityl-L-Cysteine) (Skoufias et al., 2006) during prometaphase at a point when the spindle had formed, and the kinetochores of polar chromosomes approached the main spindle surface but had not yet attached to it. After the addition of STLC, the spindle started shortening, and the kinetochores of the polar chromosomes that were already close to the spindle surface started pivoting towards the back of the spindle (Figure 3B, Video 5). This reversed pivoting was evident as an increase in the pivoting angle during spindle shortening in STLC-treated cells, in contrast to the decrease of the pivoting angle during spindle elongation in untreated cells (Figure 3E-F, Supplementary Figure 3A-C). Remarkably, the pivoting direction was reversible, moving back and forth (Figure 3D, Supplementary Figure 3D, Video 6). The washout of the Eg5 inhibitor monastrol (Mayer et al., 1999) reversed the inhibitor-induced spindle shortening into elongation, which triggered kinetochore pivoting towards the main spindle surface (Figure 3G-H). The number of kinetochore pairs behind the pole decreased before monastrol addition, increased during monastrol treatment, and decreased again after monastrol washout (Figure 3I), in agreement with the predictions from the model.

To block spindle elongation, we added the ATP-competitive Eg5 inhibitor FCPT (2-(1- (4-fluorophenyl)cyclopropyl)-4-(pyridin-4-yl)thiazole) (Groen et al., 2008; Rickert et al., 2008), which induces tight binding of Eg5 onto microtubules. Subsequently, the spindle length remained constant, and the kinetochores of polar chromosomes did not pivot significantly, providing further support to the model (Figure 3C and 3E-F, Supplementary Figure 3A-B and 3E, Video 5). Finally, to exclude the possibility that the observed effects of Eg5 inhibitors were due to roles of Eg5 other than driving spindle elongation, we employed an orthogonal approach. This involved adding a low dose of nocodazole (20 nM), which induces mild spindle shortening after NEBD while preserving the attachments of chromosomes to astral microtubules (Rajendraprasad et al., 2021). In agreement with the results from Eg5 inhibition, spindle shortening caused an increase in the number of kinetochores behind the pole (Supplementary Figure 3F-H). Collectively, these data show that the centrosome movement due to spindle elongation or shortening dictates the direction of kinetochore pivoting.

To quantitatively test our model, we explored whether the extent of pivoting can be explained by the extent of spindle elongation or shortening. A strong correlation between the change in spindle length and the simultaneous change in the pivoting angle supports this notion (Figure 3J). Moreover, we calculated the expected change in the pivoting angle for a given change in the spindle length, assuming immobile chromosomes (see Methods and Supplementary Figure 3I). The resulting theoretical prediction closely matched the linear fit of the data (Figure 3J). Taken together, our results reveal spindle elongation as a necessary and sufficient driver of chromosome passage across the spindle pole, dictating both the direction and magnitude of pivoting of chromosome-carrying astral microtubules (Figure 3K).

### Polar chromosomes form complex attachments yet remain bound to the same microtubule during pivoting

Prior to establishing end-on attachments, kinetochores initially bind to microtubules laterally (Alexander and Rieder, 1991; Magidson et al., 2011; Shrestha and Draviam, 2013). To explore the types of attachments that polar kinetochores form with astral microtubules during their pivoting, and to determine if they transition between different microtubules, we imaged microtubules by STED microscopy in fixed RPE1 cells expressing CENP-A-GFP, at various stages between the end of chromosome centripetal movement and landing (Figure 4A). Surprisingly, rather than primarily observing lateral attachments, we found a wide variety of attachment types, including lateral, monotelic, and syntelic configurations, together with more complex and previously unidentified types (Figure 4A). We define them as: 1) bilateral, when two sister kinetochores attach laterally to two astral microtubules, 2) mono-lateral, when one sister kinetochore has an end-on attachment in addition to both sisters being laterally attached to the same astral microtubule, 3) mono-bilateral, as a combination of bilateral and one end-on attachment, and 4) syn-lateral, as a combination of syntelic and one lateral attachment. Mono-lateral attachments were the most frequent, followed by lateral and bilateral ones (Figure 4B).

**Figure 4.**
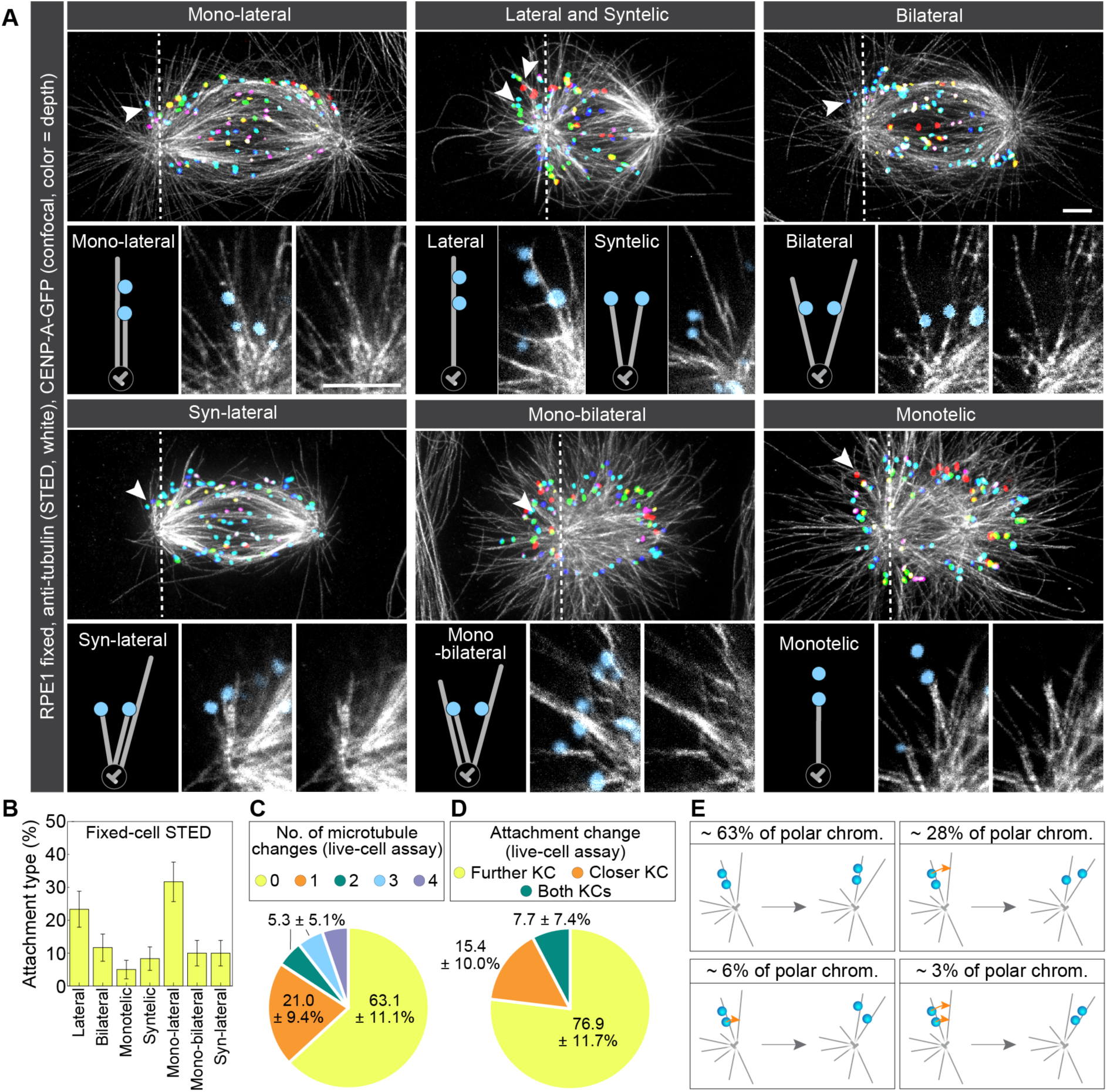
Polar kinetochores form complex attachments with the microtubules they remain attached to during pivoting. **(A)** STED superresolution images of microtubules immunostained for α-tubulin (white) in RPE1 cells stably expressing CENP-A-GFP (colorful, confocal). Images show maximum intensity projections of the whole cell. Kinetochores are color-coded for depth with the brgbcmyw LUT in ImageJ throughout 8-17 z-planes, corresponding to 2.4-5.1 µm. White arrows represent a kinetochore pair attached to microtubules shown in insets below the image. Insets show single z-planes or maximum intensity projections of 2-3 z-planes where the microtubule of interest is located, smoothed with 0.75-mm-sigma Gaussian blur. The schematic representations of different attachment types are shown next to the associated insets. **(B)** Fractions of different attachment types for polar chromosomes between the end of the centripetal movement and landing quantified from STED images. N = 60 kinetochore pairs in 24 cells from 5 experiments. **(C)** Pie chart for the number of microtubule changes during pivoting measured in live RPE1 cells stably expressing EYFP-tubulin with added HaloTag-CENP-A (See Supplementary Figure 4). N = 19 kinetochore pairs in 10 cells from 4 experiments. **(D)** Pie chart for attachment changes on further (yellow), closer (blue) and both (green) polar kinetochores measured in the same cells as in (C). N = 13 kinetochore pairs in 5 cells from 3 experiments. **(E)** Schematic representations of types of attachment changes that polar kinetochores (blue circles) undergo during pivoting and their frequencies. The orange arrows denote which kinetochore in a pair changes microtubules during pivoting. All scale bars, 2 µm.

To explore if chromosomes can switch from one microtubule to another during pivoting, we imaged live cells expressing EYFP-α-tubulin and H2B-mRFP with HaloTag-CENP-A (Supplementary Figure 4B). We found that 63.1 ± 11.1 % of polar chromosomes remained attached to a single microtubule during pivoting, 21.1 ± 9.4 % made one change between microtubules, whereas multiple changes were rare (Figure 4C). Most of the observed changes included only a change for the further kinetochore, whereas the closer one remained attached to the same microtubule during the entire process (Figure 4D and Supplementary Figure 4B). Thus, in a typical case with 7 polar chromosomes, 4 of them remain attached to a single microtubule during pivoting, 2 change the attachment of the further kinetochore, e.g. from lateral to bilateral, whereas 1 shows a more complex transition (Figure 4E). In sum, polar chromosomes make a variety of attachments with astral microtubules but typically remain attached to the same microtubules while crossing the polar region.

### Kinetochores on polar chromosomes show molecular signatures of lateral and immature end-on attachments

To characterize the molecular nature of polar kinetochore-microtubule attachments together with STED imaging of microtubules during intact early mitosis, we tested the levels of Mad2, a checkpoint protein that marks kinetochores with missing or unstable end-on attachments (Kuhn and Dumont, 2017; Waters et al., 1998). Among sister kinetochores, the one positioned further from the pole contained higher levels of Mad2 (Figure 5A-B and Supplementary Figure 4A). This suggests that the further kinetochore typically forms a lateral attachment, whereas the closer one forms an end-on attachment in addition to the lateral one, in agreement with microtubule localization observed by STED microscopy (Figure 5A). Mad2 levels at kinetochores with end-on attachments (Figure 5A) were lower than at those with lateral attachments, as expected, but higher than at end-on attached kinetochores in metaphase (Figure 5C), suggesting that the end-on attachments on polar chromosomes are immature.

**Figure 5.**
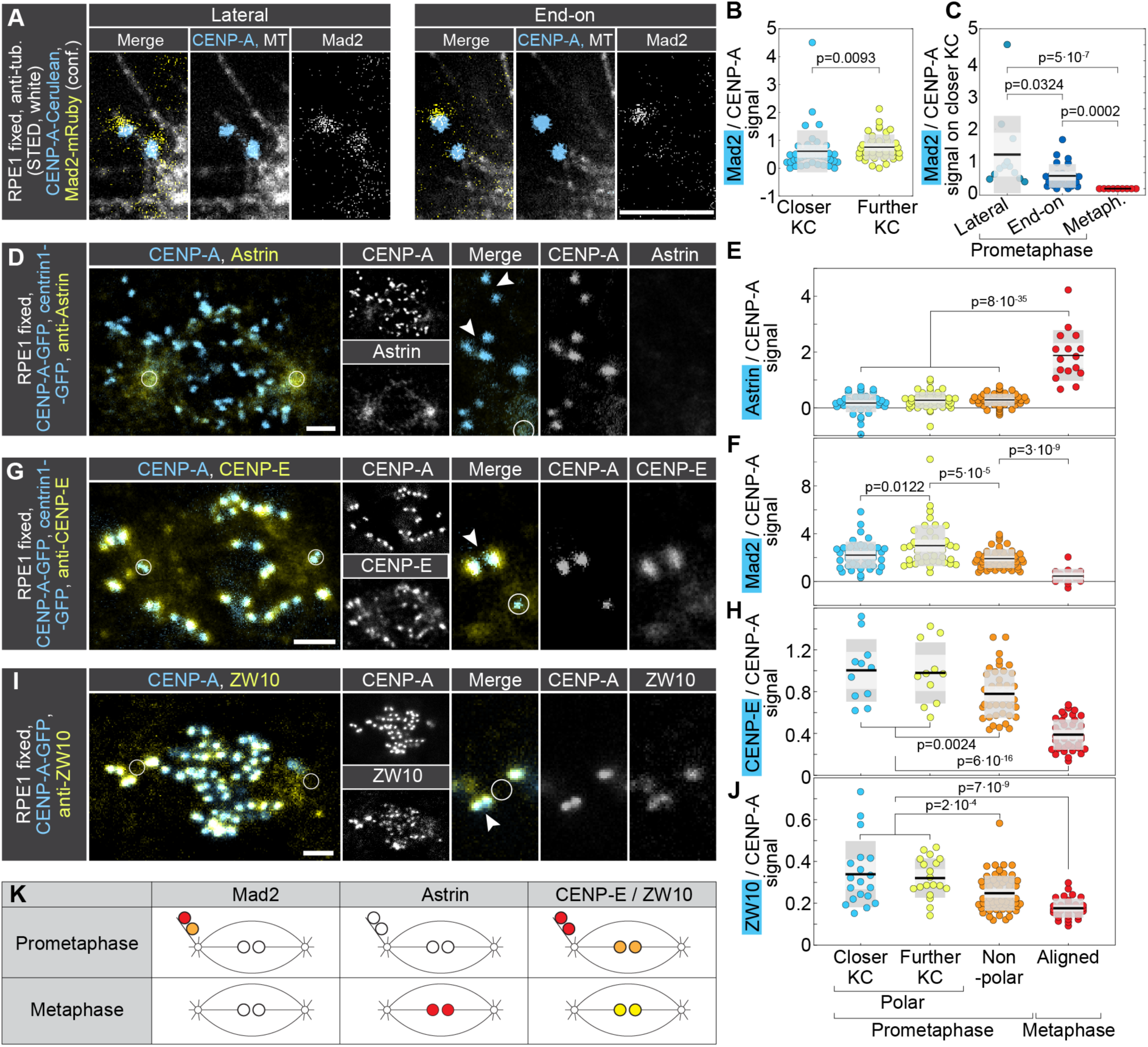
Kinetochores on polar chromosomes in early intact mitosis bind Mad2, CENP-E, and ZW10, but not Astrin. **(A)** STED superresolution images of polar chromosomes attached to microtubules immunostained for α-tubulin (white) in RPE1 cells stably expressing CENP-A-Cerulean (blue) and Mad2-mRuby (yellow). Images show a single z-plane, with kinetochores and microtubules smoothed with 0.75-mm-sigma Gaussian blur. Intensities of CENP-A and Mad2 are equally adjusted throughout the images. **(B)** Univariate scatter plot of the relative Mad2/CENP-A signal for closer (blue) and further (yellow) polar kinetochore. Measurements were done on STED images as from (A). N = 42 kinetochore pairs in 29 cells from 3 experiments. Statistical test: Mann-Whitney U test. **(C)** Univariate scatter plot of the relative Mad2/CENP-A signal for closer polar kinetochore in metaphase (blue), in prometaphase when the kinetochore makes an end-on attachment (orange) and in prometaphase when the kinetochore makes a lateral attachment (yellow). Measurements are done on STED images as from (A). N = 10 kinetochores from 5 kinetochore pairs in 2 cells from 1 experiment for metaphase, N = 26 end-on attached kinetochores and 12 laterally attached kinetochores in 29 cells from 3 experiments for prometaphase. Statistical test: Kruskal-Wallis with post-hoc Dunn’s test. **(D–I)** Confocal images of mitotic spindles in early prometaphase cells expressing CENP-A-GFP and centrin-1-GFP (cyan), (D, G), or CENP-A-GFP alone (cyan, I), immunostained for Astrin (yellow, D) and Mad2 (not shown, D), CENP-E (yellow, G), and ZW10 (yellow, I). Individual fluorescence channels are shown on the right, along with merged enlargements of the polar region and the corresponding individual channels. Circles denote centrosome position and arrowheads point to polar kinetochores. Quantification of mean Astrin (E), Mad2 (F), CENP-E (H), and ZW10 (J) signal intensities, normalized to mean CENP-A levels at kinetochores, is shown for the specified spindle positions and mitotic phases. N = 116 kinetochore pairs in 20 cells (E and F) from 3 experiments; 92 kinetochore pairs in 8 cells (H), and 115 kinetochore pairs in 10 cells (J), from 2 experiments. Statistical test in (E), (F), (H), (J): ANOVA with post-hoc Tukey test. **(K)** Scheme of the observed protein levels (red: high, orange: medium, yellow: low, white: absent) on polar and non-polar kinetochores (circles to the upper left of the spindle and in the spindle center, respectively) in prometaphase and metaphase, as indicated. All scale bars, 2 µm.

To assess a positive marker of end-on attachments, we examined Astrin, which localizes to aligned kinetochores with low Aurora B activity, but is absent from unaligned kinetochores (Mack and Compton, 2001; Schmidt et al., 2010). Consistent with these findings, we observed that Astrin was absent from both polar kinetochores within a pair irrespective of their relative distance to the centrosome (Figure 5D-E). It was also absent from non-polar kinetochores in prometaphase, which had intermediate levels of Mad2, but was present on the kinetochores aligned at the metaphase plate, which were devoid of Mad2 (Figure 5E-F).

The levels of CENP-E and ZW10, components of the fibrous corona that promotes lateral attachments (Yao et al., 1997), were high at polar kinetochores, including both sisters, indicating the presence of a large corona (Figure 5G-J). In comparison with polar kinetochores, non-polar kinetochores on prometaphase spindles recruited less CENP-E and ZW10, whereas the lowest signal was at aligned kinetochores in metaphase (Figure 5G-J). Taken together, these results suggest that in early intact mitosis, both sister kinetochores on polar chromosomes typically form lateral attachments, while the sister closer to the pole additionally establishes immature end-on attachments characterized by the presence of checkpoint and corona proteins and the absence of Astrin. These attachments mature as the kinetochores interact with the spindle body and congress, recruiting Astrin and losing corona and checkpoint proteins (Vukušić and Tolić, 2024) (Figure 5K).

### Microtubules from the opposite spindle half help the movement of polar chromosomes near the spindle surface

To explore the contribution of microtubules extending from the other spindle half to the movement of polar chromosomes (Figure 1E), we used STED microscopy and found that in phases between the end of centripetal movement and landing, 30.0 ± 5.9 % of kinetochore pairs in the polar region (i.e., those attached to an astral microtubule and within ∼3 μm from the spindle pole) were located next to a microtubule extending from the opposite side of the spindle (Figure 6A), whereas 70.0 ± 5.9 % had no such microtubule nearby (Figure 6B). Attachments to microtubules from the opposite spindle half were rare behind the pole, seen in only 20.5 ± 6.5 % kinetochore pairs, but were more frequent in the region in front of the pole, where 47.6 ± 10.9 % of kinetochore pairs showed attachments before joining the main spindle surface (Figure 6B). Although we could not determine whether these microtubules originated at the opposite pole or along the microtubules of the opposite spindle half, their origin at the opposite side indicates that their plus end was facing the kinetochore. In agreement with this orientation, these microtubules were attached to kinetochores predominantly in an end-on configuration (Figure 6C, Supplementary Figure 5A). Most of these microtubules had tubulin intensity similar to that of astral microtubules, suggesting they are single microtubules (Supplementary Figure 5B).

**Figure 6.**
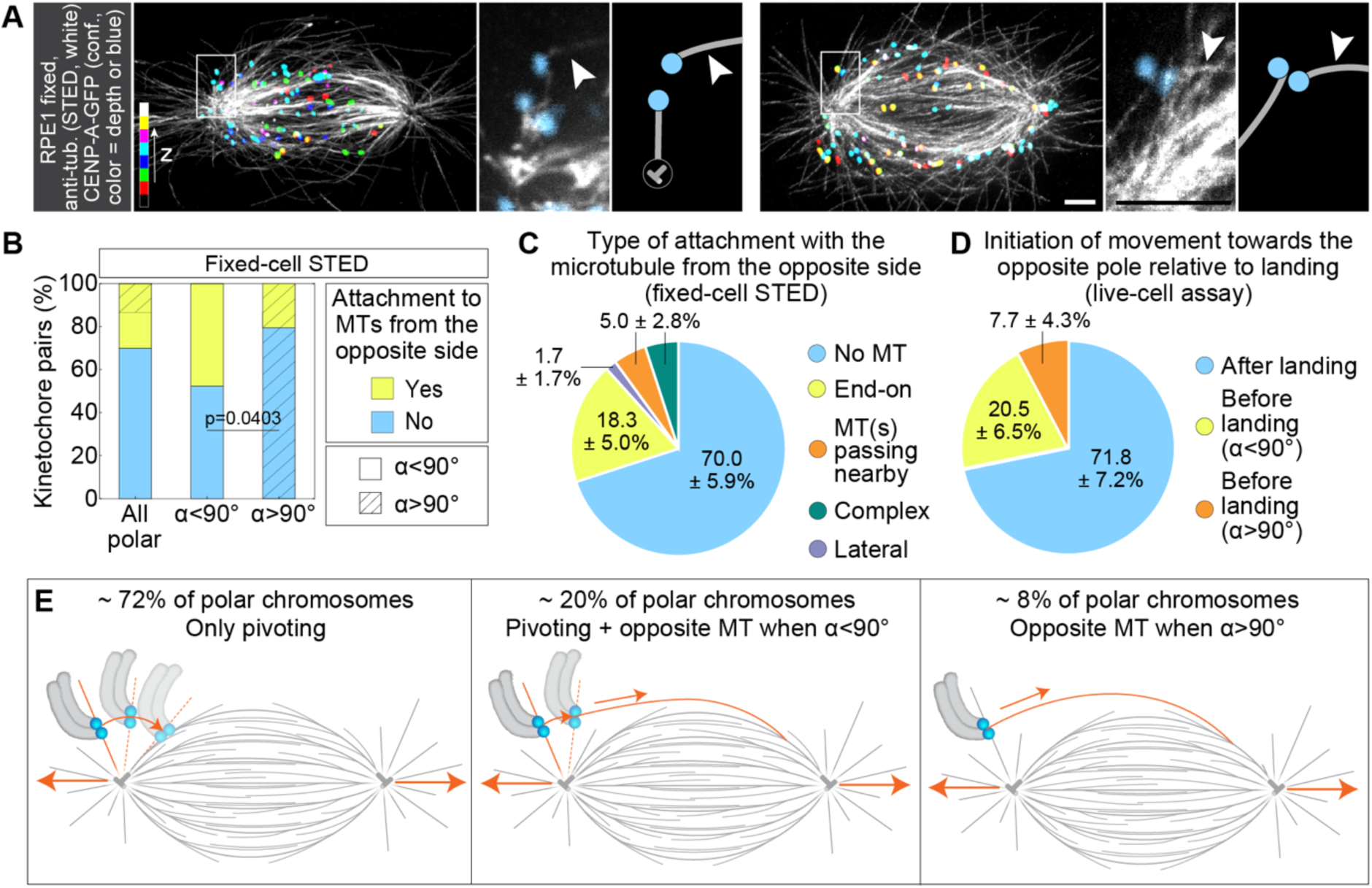
Microtubules from the opposite side of the spindle facilitate the movement of polar chromosomes in the region close to the spindle surface. **(A)** STED superresolution images of microtubules immunostained for α-tubulin (white) in RPE1 cells stably expressing CENP-A-GFP (colorful, confocal). Images show maximum intensity projections of the whole cell. Kinetochores are color-coded for depth with the brgbcmyw LUT in ImageJ throughout 16 and 9 z-planes, corresponding to 4.8 and 2.7 µm, respectively. White boxes represent regions containing polar kinetochores attached to the microtubule from the opposite side of the spindle, which are enlarged to the right and represented schematically. White arrows denote the microtubule coming from the opposite side of the spindle. Enlargements are maximum intensity projections of 2 and 3 z-planes where the microtubule of interest is located, smoothed with 0.75-mm-sigma Gaussian blur. **(B)** Percentages of kinetochores that are unattached (blue) or attached to the microtubule from the opposite side of the spindle (yellow) measured in cells as in (A). The shaded area represents kinetochores located at the angle above 90° with respect to the long spindle axis. N = 18 kinetochore pairs in 15 cells from 5 experiments. Statistical test: Fisher’s exact test. **(C)** Pie chart representing the percentage of different attachment types with the microtubule from the opposite side. N = 18 kinetochore pairs in 15 cells from 5 experiments. **(D)** Pie chart representing the percentage of polar kinetochores from LLS videos that initiate their movement towards the opposite pole after landing (blue), before landing at an angle below 90° (yellow) and before landing at an angle above 90° (orange), indicative of a force other than pivoting. N = 39 kinetochore pairs in 20 cells from 8 experiments. **(E)** Schematic representation of percentages of polar chromosomes that pivot without forming an attachment to the microtubule from the opposite side before landing (left), that pivot and attach to microtubule from the opposite side once they reach the angle below 90° (middle) and the ones that attach to microtubule from the opposite side at the angle above 90° (right). Scale bars, 2 µm.

If microtubules from the opposite side pull the chromosome across the pole, this is expected to increase the distance between the kinetochores and the closer pole, as well as the interkinetochore distance, assuming that these microtubules attach to the sister kinetochore that is not end-on attached to the closer pole. We found such changes within one minute before landing (Supplementary Figure 5C-D), when the kinetochores are typically close to the main spindle surface (at an angle <60°, see Figure 2E), suggesting that the pulling by microtubules from the opposite side contributes mainly to the last part of the chromosome movement towards the spindle surface. Live-cell STED microscopy provided direct visualization of the process in which such kinetochores attain end-on attachment to a microtubule extending from the opposite spindle half, followed by an increase in interkinetochore distance and movement towards the opposite pole (Supplementary Figure 5E). These findings show that microtubules extending from the opposite side of the spindle can pull the chromosome, especially when it is already close to the spindle surface. Together, our results suggest that of the 7 polar chromosomes, around 5 cross the polar region and reach the main spindle surface only by elongation-driven pivoting, 1-2 are pulled by microtubules from the opposite side after the initial pivoting, whereas <1 chromosome per cell interacts with microtubules from the opposite side before pivoting (Figure 6E).

### Incomplete spindle elongation leads to delayed chromosome passage across the pole and missegregation

To explore the functional relevance of the pivoting model, we asked whether incomplete spindle elongation may lead to missegregation of polar chromosomes. To obtain a system with chromosome segregation errors in RPE1 cells, we inhibited the mitotic kinase Mps1, a key element of the spindle assembly checkpoint and the correction of erroneous attachments, by the small-molecule inhibitor AZ3146 (Hewitt et al., 2010) (Figure 7A, Video 7). We reasoned that due to the premature anaphase onset, spindle elongation may be incomplete in a subset of cells. This should in turn lead to incomplete passage of polar chromosomes across the pole in those cells, allowing us to explore the consequences of perturbed pole crossing on the segregation fidelity of these chromosomes.

**Figure 7.**
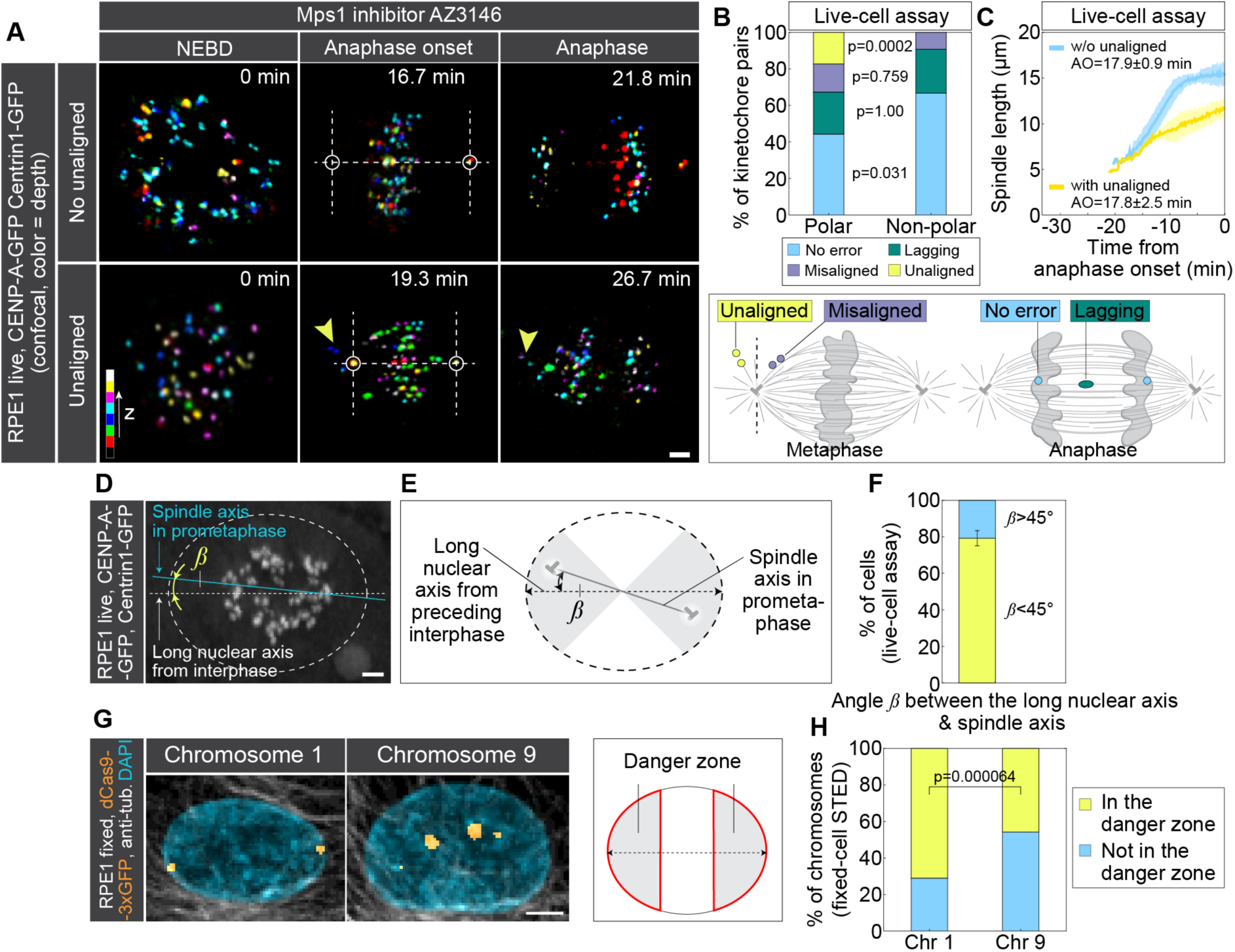
Incomplete spindle elongation compromises passage across the pole for polar chromosomes located in the danger zone. **(A)** Time-lapse images of RPE1 cells stably expressing CENP-A-GFP and Centrin1-GFP (colorful, confocal) and treated with Mps1 inhibitor AZ3146. The images show maximum intensity projections of the imaged cells. The upper panel represents cells without unaligned chromosomes and the lower panel represents cells with unaligned chromosomes. Kinetochores are color-coded for depth with the brgbcmyw LUT in ImageJ throughout 14-21 z-planes, corresponding to 5.6-8.4 µm. **(B)** Percentage of polar (left bar) or non-polar (right bar) kinetochore pairs, as determined at NEBD, that segregated normally (no error, light blue) or ended up as lagging (green), misaligned (dark blue) or unaligned (yellow) kinetochores in anaphase. Each group of kinetochores is depicted on the scheme below the graph. N = 52 polar kinetochore pairs in 7 cells from 4 experiments. N = 54 non-polar kinetochore pairs in 7 cells from 4 experiments. Statistical test: Fisher’s exact test. **(C)** Spindle length in time for cells treated with AZ3146 divided into those that have (orange) or do not have (blue) unaligned chromosomes at anaphase onset. Values are shown as mean (dark line) and SEM (shaded areas). N = 3 cells with unaligned chromosomes from 2 experiments. N = 4 cells without unaligned chromosomes from 2 experiments. **(D)** RPE1 cell imaged using lattice light-sheet microscopy at the end of centripetal movement. The dashed oval denotes the approximate position of the nucleus in the preceding interphase (see Supplementary Figure 6). The straight white line denotes the position of the long nuclear axis in the preceding interphase, the blue line is the spindle axis in prometaphase at the end of the centripetal movement, and β is the angle between them. The image is a maximum intensity projection of 42 z-planes covering the whole cell, corresponding to 6.1 µm. **(E)** Scheme showing the angle β as in (D), grey areas denote β < 45°. **(F)** Percentage of cells with β < 45° (yellow) and β > 45° (blue). N = 24 cells from 8 experiments. **(G)** Images show nuclei in interphase in fixed RPE1 cells stably expressing dCas9-3xGFP (orange) and stained with anti-α-tubulin (gray) and DAPI (blue). Cells are transduced with lentiviruses for chromosome 1 or chromosome 9, as indicated. Images are maximum intensity projections of 30 z-planes, corresponding to 15 µm. The diagram on the right illustrates the "danger zone," which consists of two nuclear caps. Each cap covers one-third of the nuclear long axis. Chromosomes positioned in these caps during interphase are more likely to end up behind the centrosome in early mitosis. **(H)** Percentage of cells with chromosome 1 or 9 located in the nuclear danger zone (yellow) or outside the nuclear danger zone (blue). N = 266 chromosomes in 133 cells from 4 experiments for chromosome 1. N = 72 chromosomes in 36 cells from 3 experiments for chromosome 9. Statistical test: Chi-squared test. All scale bar, 2 µm.

The chromosomes that were polar at NEBD missegregated more frequently than non-polar chromosomes that were similarly positioned with respect to the nucleus and the centrosome midpoint in Mps1-inhibited cells (Figure 7B) (Klaasen et al., 2022). Whereas polar and non-polar chromosomes had a similar tendency to end up as misaligned and lagging chromosomes in anaphase, there was a stark difference in the occurrence of unaligned chromosomes (Figure 7B). About 20% of polar chromosomes remained unaligned throughout mitosis, leading to aneuploid daughter cells. In contrast, such cases were absent in the non-polar group (Figure 7B, Video 7). These results suggest that the missegregation bias towards polar chromosomes is largely caused by their persistent unalignment.

Among the chromosomes with erroneous segregation, the persistently unaligned chromosomes were the last ones to pass from the back to the front of the spindle pole (Supplementary Figure 6A), suggesting that the cause for their unalignment lies in perturbed passage around the pole. Based on the pivoting model, we expect that the cells that entered anaphase with persistently unaligned chromosomes had less extensive spindle elongation than those without unaligned chromosomes. Indeed, we found that among the Mps1-inhbited spindles, those that had persistently unaligned chromosomes elongated more slowly, entering anaphase at a ∼4 μm shorter length than the spindles without such chromosomes (Figure 7C, Supplementary Figure 6B-C). Different timing of anaphase was not the cause of persistent unalignment, as the spindles with and without unaligned chromosomes entered anaphase at a similar time (17.8 ± 2.5 vs. 17.9 ± 0.9 minutes). These data show that the propensity of polar chromosomes to remain unaligned is higher in cells with slower spindle elongation, suggesting the importance of effective chromosome passage across the pole.

The pivoting model may be linked with chromosome identity, as certain chromosomes missegregate more frequently than others across various treatments and cancer types (Klaasen et al., 2022; Worrall et al., 2018). We hypothesize that there is a zone within the interphase nucleus, related to the axis of the future spindle, that would predispose chromosomes to missegregate as persistently unaligned, which we term “danger zone.” In most RPE1 cells, centrosome separation occurs before NEBD, positioning one centrosome above and the other below the nucleus, followed by spindle elongation and rotation that aligns the spindle parallel to the coverslip surface (Magidson et al., 2011). Within this plane, we observed that in prometaphase, the centrosome-centrosome axis aligns closely with the long nuclear axis from the preceding interphase, as reflected by an angle of 26.1° ± 4.2° between the two axes (N = 24 cells, Figure 7D-F, Supplementary Figure 6D-E, Video 8). Furthermore, the nuclear axis in interphase is typically aligned with the long cellular axis (Supplementary Figure 6F, Video 8) (Stiff et al., 2020). This suggests that the danger zone is situated in the nuclear caps, defined as regions near the ends of the major axis of the nucleus (Figure 7G, diagram), because the chromosomes located there in interphase typically end up behind the centrosome in early mitosis.

We then compared interphase positions of chromosomes 1 and 9 with respect to the identified danger zone, which show a missegregation bias and random missegregation, respectively (Klaasen et al., 2022) (Figure 7G). Remarkably, we found that chromosome 1 resides in the danger zone in 71.00 ± 2.78 % of cases. In contrast, chromosome 9 resides in the danger zone in only 45.83 ± 5.87 % of cases (Figure 7H). These findings support the idea that the missegregation bias of chromosome 1 arises from its preferential position in interphase within the nuclear caps. Chromosomes located in this danger zone are susceptible to failure in crossing the pole, leading to their persistent unalignment during mitosis (Klaasen et al., 2022).

### Spindle elongation helps to resolve polar chromosomes in cancer cell lines

Finally, we set out to investigate the universality of the pivoting model by testing cancer-derived cell lines. To determine whether microtubules attached to polar chromosomes pivot around the spindle pole in cancer cells, we used the osteosarcoma U2OS cell line expressing GFP-tagged EB3 (Stepanova et al., 2003) and CENP-A-mCherry. We visualized microtubule contours using time projections of EB3 spots and observed that microtubules associated with polar kinetochores pivoted around the pole as it moved outward during spindle elongation (Figure 8A, Video 9). Tracking of kinetochores from NEBD to landing in live U2OS cells expressing CENP-A-GFP and tubulin-mCherry showed that polar kinetochores transitioned from the back to the front of the spindle pole, reflected by a decrease in their pivoting angle, during spindle elongation (Figure 8B-D, Supplementary Figure 7A-B, Video 10). These findings indicate that microtubule pivoting occurs during spindle elongation in U2OS cells similarly to RPE1 cells.

**Figure 8.**
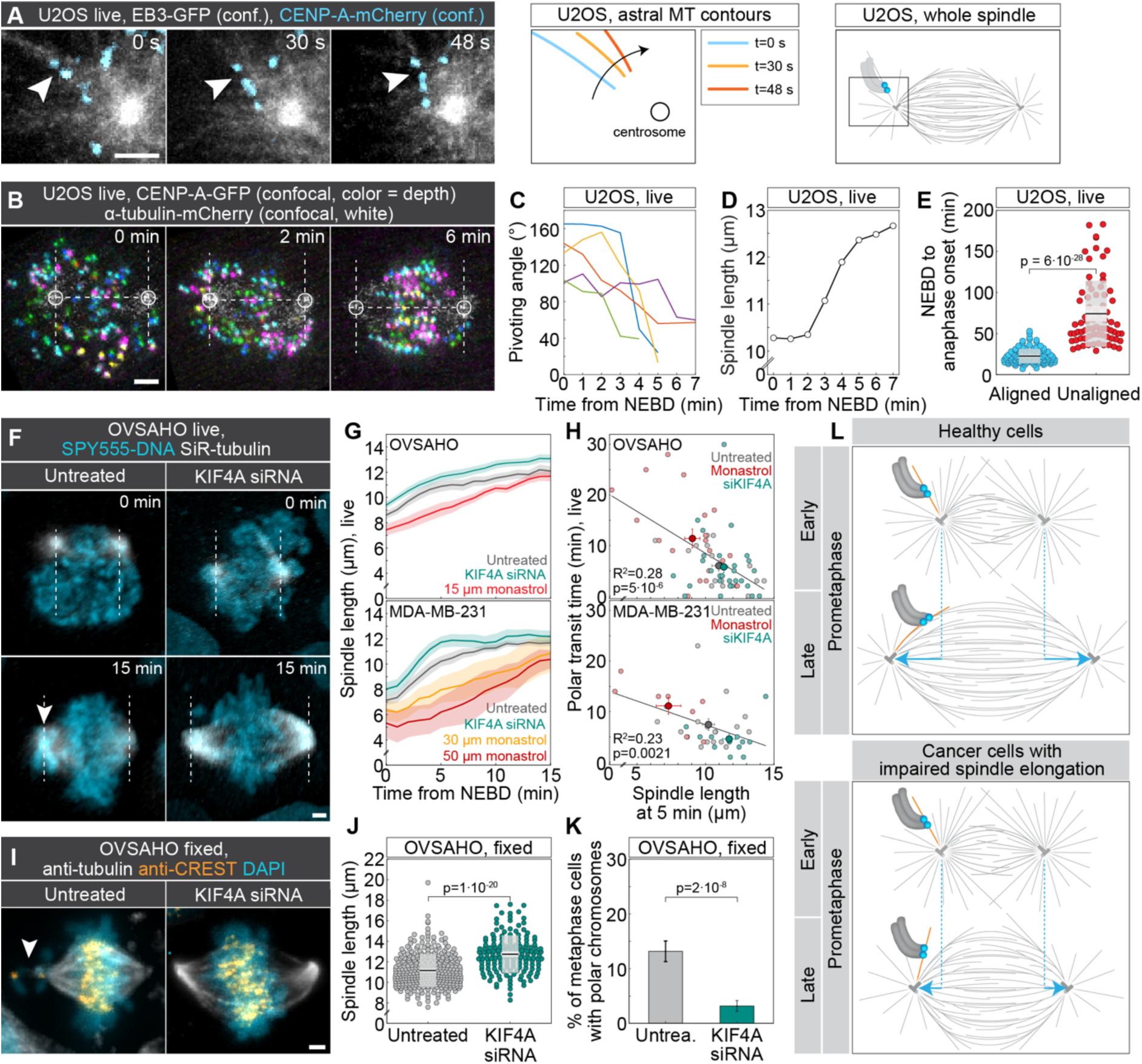
Spindle elongation promotes chromosome alignment in cancer cell lines. **(A)** Time-lapse confocal images of live U2OS cells stably expressing EB3-GFP (white, confocal) and CENP-A-mCherry (blue, confocal) showing astral microtubule attached to the polar kinetochore pair and pivoting over time. Images are temporal maximum intensity projections of a single z-plane in 3-5 sequential time frames during which the EB3 spot was visible. Astral microtubule contours during the pivoting are shown to the right, and a scheme showing the imaged region within the spindle is at the far right. (**B**) Time-lapse confocal images of live U2OS cells stably expressing CENP-A-GFP (colored for depth) and tubulin mCherry (white). Circles mark the poles and dashed lines the spindle axis and the edge of the polar region. (**C-D**) Pivoting angle for 5 polar kinetochore pairs (C) and spindle length (D) over time for the cell shown in (B). (**E**) Univariate scatter plot of the time from NEBD until anaphase onset in live U2OS cells as in (B), comparing cells with aligned chromosomes (left, N = 111) and those that had one or more chromosomes that were not aligned (right, N = 73) at the end of spindle elongation. Statistical test: t-test. **(F)** Time-lapse images of control (left) and Kif4A siRNA-treated (right) live OVSAHO cells stained with SPY-555-DNA (blue) and SiR-tubulin (white), imaged using lattice light-sheet microscopy. Dashed lines denote the edge of the polar regions; arrowhead points to chromosomes near the pole. **(G)** Spindle length over time for OVSAHO (top; N = 19, 22, 20 cells for the indicated treatments) and MDA-MB-231 cells (bottom; N= 18, 22, 5, 3 cells for the indicated treatments) treated as stated in the legend. Lines, mean; shaded areas, SEM. (**H**) Polar transit time for OVASHO (top) and MDA-MB-231 cells (bottom) as a function of the spindle length measured 5 minutes after NEBD. Small circles show individual cells, and large circles with error bars are mean and SEM for each treatment. Linear regression on individual data points is also shown. **(I)** Images of fixed OVSAHO cells, control (left) and KIF4A siRNA-treated (right), stained with anti-α-tubulin antibody (white), anti-CREST (orange), and DAPI (blue). The white arrow marks a chromosome at the pole in metaphase. **(J)** Univariate scatter plot of the spindle length in untreated (N = 319) and KIF4A siRNA-treated (N = 157) fixed OVSAHO cells. Statistical test: t-test. (**K**) Percentage of metaphase cells with polar chromosomes in untreated and KIF4A siRNA-treated fixed OVSAHO cells from (J). Statistical test: Chi-squared test. In (B), (F), and (I), images are maximum intensity projections of the whole cell. All scale bars, 2 µm. **(L)** A model of how polar chromosomes pass across the pole in normal cells and fail to cross the pole when spindle elongation is impaired, such as in certain cancer cells. In healthy cells, polar chromosomes are attached to astral microtubules. While the chromosomes remain largely immobile, the spindle elongates, thereby driving the pivoting of astral microtubules around the spindle pole. Consequently, the chromosomes with their microtubules get close to the main spindle surface, where they attach and move to the spindle midplane. In cells with impaired spindle elongation, this elongation is insufficient to drive microtubule pivoting from the back of the spindle pole all the way towards the main spindle surface, leaving polar chromosomes persistently unaligned.

To evaluate the efficiency of the pivoting mechanism in this U2OS cell line, which has chromosome congression defects (Klaasen et al., 2022), we tracked and counted polar kinetochores over time (Methods). We found that 96.1% of polar kinetochores successfully crossed the pole and congressed to the metaphase plate, whereas 3.9% remained unaligned, i.e., behind the pole, at the end of spindle elongation (N = 44 out of 1127 polar kinetochore pairs from 101 cells). These results suggest that the pivoting mechanism is highly efficient in U2OS cells and facilitates successful congression of most polar kinetochores. Tracking of kinetochores that remained unaligned revealed that they eventually moved to the main spindle surface (Supplementary Figure 7C), likely through capture by microtubules from the opposite spindle half. However, this elongation-independent movement was significantly delayed, as indicated by a threefold increase in mitotic duration in cells retaining an unaligned chromosome after spindle elongation (Figure 8E). Thus, failure of timely polar chromosome transit during spindle elongation prolongs mitosis in a cancer cell line.

According to our model, slower spindle elongation should delay the incorporation of polar chromosomes into the spindle body, whereas enhanced elongation should promote it. To test this prediction, we perturbed spindle elongation in two highly chromosomally unstable cancer cell lines, OVSAHO (ovarian carcinoma) and MDA-MB-231 (triple-negative breast cancer), both characterized by congression defects (Tamura et al., 2020; Tucker et al., 2023) (Supplementary Figure 7D). We used low-dose monastrol to slow spindle elongation and KIF4A depletion to enhance it (Jagrić et al., 2021) (Figure 8F, Supplementary Figure 7E-G, Video 11 and 12). Monastrol concentrations were titrated in each line to reduce spindle elongation post-NEBD without inducing monopolar arrest in most cells (Methods). Live-cell imaging confirmed that monastrol-treated spindles were shorter and KIF4A-depleted spindles were longer than untreated ones throughout early mitosis in OVSAHO and MDA-MB-231 cells (Figure 8G). The duration of polar chromosome transit, defined as the time when the last polar chromosome moved from the rear to the front side of the spindle pole, was inversely correlated with spindle length in early prometaphase in both cell lines (Figure 8H), in agreement with our model.

To extend our findings to larger cell populations, we analyzed fixed, immunostained metaphase cells treated with either KIF4A siRNA to increase spindle length or low-dose nocodazole (10 ng/mL) to reduce spindle length (Supplementary Figure 7H). Nocodazole was selected over monastrol for the fixed-cell assay, as monastrol-treated spindles reached near-normal length by metaphase (Figure 8G). In addition to OVSAHO and MDA-MB-231, we analyzed the Caco-2 cell line (colorectal adenocarcinoma) (Thompson and Compton, 2008). These experiments revealed an overall trend where cells with longer spindles had fewer polar chromosomes than those with shorter spindles (Supplementary Figure 7I). In OVSAHO cells, KIF4A depletion increased spindle length from 11.16 ± 0.09 μm (N = 319) to 12.74 ± 0.15 μm (N = 157), and this was associated with a fourfold reduction in the fraction of cells harboring polar chromosomes (from 13.2 ± 2.1% to 3.2 ± 1.4%) (Figure 8I-K). The distance of polar chromosomes from the nearest pole was comparable between untreated and KIF4A-depleted cells (Supplementary Figure 7J), indicating that polar ejection forces (Chong et al., 2024; Novais-Cruz et al., 2023) were not significantly altered. Conversely, nocodazole treatment shortened spindle length and increased the number of polar chromosomes relative to controls (Supplementary Figure 7I), likely due to both spindle shortening and microtubule stabilization (Rajendraprasad et al., 2021). MDA-MB-231 and Caco-2 cells exhibited changes in the same direction as OVSAHO following KIF4A depletion and nocodazole treatment, though the effect of Kif4A was not statistically significant (Supplementary Figure 7I).

In conclusion, our results demonstrate that the spindle elongation-driven pivoting model applies across multiple cancer cell lines and plays a key role in ensuring rapid mitosis. Treatments that reduce spindle elongation impair the resolution of polar chromosomes, whereas enhanced elongation facilitates their timely incorporation into the spindle.

## Discussion

### The pivoting model

Our study uncovers an essential step in chromosome congression that safeguards genome integrity by rescuing polar chromosomes, which are at the highest risk of missegregation (Klaasen et al., 2022), from the “danger zone” behind the spindle pole. During early prometaphase, once the centripetal movement of chromosomes ends, the spindle begins to elongate (Magidson et al., 2011). During this phase, the poles push outward, moving beyond the polar chromosomes, which are relatively stationary. As the chromosomes remain attached to astral microtubules throughout his process, these microtubules passively pivot around the spindle pole from the outer to the inner spindle region, ultimately reaching the spindle surface and enabling the interaction of polar chromosomes with it and subsequent congression (Figure 8L).

Our findings indicate that, once the centripetal chromosome movement ends, polar chromosomes usually remain attached to the same microtubule bundle extending from the centrosome until approaching the spindle surface. If the microtubules were to extend at a constant angle from the centrosome without the ability to pivot, the chromosome would move along with the centrosome, away from the spindle center. However, instead of moving, the chromosome remains stationary, and it is the angle of the microtubules that changes, repositioning the chromosome from behind the centrosome to its front side where it can congress to the spindle midplane. Taking an evolutionary perspective, since a similar type of microtubule rotational movement has been observed in yeast (Kalinina et al., 2013), it will be valuable to assess how conserved microtubule pivoting, its underlying mechanisms, and functions are across various species.

The mechanism we propose relies on a biomechanical property of the microtubule-centrosome connection that enables the microtubules to pivot under force. Given that the pericentriolar material anchors the microtubules at the centrosome (Woodruff et al., 2014), it will be interesting to explore how mechanical properties of the pericentriolar material affect microtubule pivoting and, in turn, the rescue of polar chromosomes. Remarkably, the old and the young centrosomes behave differently in mitosis and the old one retains more unaligned chromosomes (Gasic et al., 2015). It may be that microtubule pivoting is different on the old and young centrosome and that this leads to a bias in chromosome alignment errors.

### The role of microtubule pivoting in spindle assembly

The pivoting of microtubules attached to polar chromosomes may play a role in spindle assembly. We found that typical polar chromosomes arrive at the spindle surface with mono-lateral attachments. The microtubules in the “mono” part of the attachment may represent an early kinetochore fiber, which, upon chromosome arrival to the spindle surface, becomes incorporated into the growing spindle. Thus, kinetochore fibers of polar chromosomes may start to form already when the chromosome is behind the pole, which is much earlier than previously thought and can explain the observed fast chromosome biorientation occurring as soon as the chromosome interacts with the spindle surface (Renda et al., 2022; Vukušić and Tolić, 2024).

We propose that microtubules laterally attached to the polar chromosome also integrate into the spindle, forming antiparallel overlaps with microtubules extending from the opposite side. This antiparallel bundle likely facilitates the formation of end-on attachments on the sister kinetochore facing the further pole by guiding the growth of new microtubules towards this kinetochore. As end-on attachments of both sister kinetochores mature and lateral attachments diminish (Magidson et al., 2011), the previously laterally attached microtubules may turn into bridging fibers that connect sister kinetochore fibers (Kajtez et al., 2016; Simunić and Tolić, 2016). Thus, polar chromosomes bring along components of their fibers to the growing spindle.

### The role of spindle elongation in maintaining genomic stability

Our findings show that incomplete or slow spindle elongation, accompanied by reduced microtubule pivoting, can lead to chromosome missegregation due to the delayed or unsuccessful arrival of polar chromosomes to the spindle surface. Previous work has shown that HeLa cells following the prophase pathway of spindle assembly, where centrosomes quickly separate to the opposite sides of the nucleus before NEBD, exhibit fewer chromosome segregation errors than those following the prometaphase pathway, where centrosome separation is slower and continues during prometaphase (Kaseda et al., 2011). Our model suggests that the differences in segregation errors between the two pathways arise from the presence of polar chromosomes (Vukušić and Tolić, 2022). In the prophase pathway, rapid spindle elongation ensures that no chromosomes are left behind the pole at NEBD, resulting in error-free segregation. In contrast, the slower centrosome separation in the prometaphase pathway leads to the presence of polar chromosomes, whose delayed arrival at the metaphase plate may increase the likelihood of segregation errors.

The spatial arrangement of chromosomes and centrosomes at the start of mitosis may explain why certain chromosomes are more prone to missegregation (Klaasen et al., 2022). As the chromosomes that are found behind the spindle pole in early mitosis are typically positioned at the nuclear periphery during the preceding interphase (Klaasen et al., 2022), our proposed model identifies the biomechanical principles underlying the biased missegregation of peripheral chromosomes.

In the context of disease, we demonstrate that insufficient spindle elongation, leading to impaired microtubule pivoting, delays the alignment of polar chromosomes in several cancer cell lines. Conversely, enhancing spindle elongation accelerates polar chromosome alignment. However, additional mechanisms may contribute to efficient chromosome alignment. For instance, in cell lines with short spindles but minimal accumulation of polar chromosomes, such as the ovarian cancer cell line OVKATE (Risteski et al., 2024; Tamura et al., 2020), pulling forces exerted by microtubules from the opposite spindle half may facilitate the integration of polar chromosomes into the spindle. The relative contribution of these alternative mechanisms across different cell lines and over time remains an interesting question for future investigation. We further propose that the failure of polar chromosomes to cross the pole during spindle elongation, observed in a subset of cancer cells and after nocodazole treatment, may be linked to increased stabilization of end-on attached microtubules (Bakhoum et al., 2009; Rajendraprasad et al., 2021; Vasquez et al., 1997), giving rise to thick microtubule bundles with limited pivoting capacity. The resulting delay in the movement of polar chromosomes may contribute to prolonged mitosis in cancer cell lines with chromosome congression defects.

We speculate that the spindle elongation-driven microtubule pivoting may be relevant to explain and modify errors in chromosome alignment in cancer cells. This idea is supported by the fact that cells without PTEN, the second most frequently mutated tumor suppressor after p53 (Li et al., 1997; Turnham et al., 2020; Yin and Shen, 2008), recruit less Eg5 to their spindles, resulting in both shorter spindles and misaligned chromosomes (He et al., 2016). Thus, a PTEN mutation might be one of the indicators that a particular type of cancer has unaligned chromosomes that stem from impaired spindle elongation and might be rescued by restoring PTEN function. More broadly, we propose that manipulating spindle elongation in certain cancer cells could be used to influence mitotic errors, either correcting them to maintain genome stability or worsening them to a degree that compromises cell viability.

## Materials & Methods

### Cell lines and plasmids

The experiments were performed using the following cell lines: human hTERT-RPE1 (hTERT-immortalized retinal pigment epithelium) cells stably expressing CENP-A-GFP, human hTERT-RPE1 (hTERT-immortalized retinal pigment epithelium) cells stably expressing CENP-A-GFP and Centrin1-GFP (Magidson et al., 2011), and human hTERT-RPE1 cells stably expressing CENP-A-GFP and Mis12-mCherry, a gift from Alexey Khodjakov (Wadsworth Center, New York State Department of Health, Albany, NY, USA); human hTERT-RPE1 cells stably expressing CENP-A-Cerulean and Mad2-mRuby (Marcozzi and Pines, 2018), a gift from Jonathon Pines (Institute of Cancer Research, London, UK); human hTERT-RPE1 cells stably expressing tubulin-EYFP and H2B-mRFP (Holland et al., 2012), a gift from Andrew Holland (Johns Hopkins University School of Medicine, Baltimore, MD, USA); human hTERT-RPE1 cells stably expressing dCas9-GFP-3xFKBP (Truong et al., 2023), a gift from Susanne Lens (Utrecht University, Utrecht, Netherlands); human hTERT-RPE1 cells stably expressing EB3-GFP and H2B-RFP, a gift from Patrick Meraldi (University of Geneva) and generated by Will Krek; unlabeled human OVSAHO cell line (JCRB1046 Cell Bank, Tebubio, Le Perray-en-Yvelines, France); unlabeled human MDA-MB-231 cell line, a gift from Dragomira Majhen (Ruđer Bošković Institute, Zagreb, Croatia); unlabeled human Caco-2 cell line (86010202, Sigma Aldrich); and human U2OS cells stably expressing EB3-2xGFP and CENP-A-mCherry (Kajtez et al., 2016).

The plasmids used for the experiments were: pHTN HaloTag-CENP-A (Castrogiovanni et al., 2022), a gift from Patrick Meraldi (University of Geneva, Geneva, Switzerland); psPAX2 (Addgene, No. 12260); psMD2.G (Addgene, No. 12259); pLH sgRNA Chr1-telo (Dumont et al., 2020) and pLH sgRNA Chr9-cen (Truong et al., 2023), both gifts from Susanne Lens (Utrecht University, Utrecht, Netherlands).

### Cell culture

All cell lines, except an OVSAHO cell line, were cultured in Dulbecco’s Modified Eagle’s Medium (DMEM) containing 1 g/L D-glucose, pyruvate and L-glutamine (Gibco), supplemented with 10% (vol/vol) heat-inactivated Fetal Bovine Serum (FBS, Sigma Aldrich, St. Louis, MO, USA) and penicillin (100 IU/mL)/streptomycin (100 mg/mL) solution (Lonza, Basel, Switzerland). The OVSAHO cell line was maintained in RPMI 1640 medium containing L-glutamine and sodium bicarbonate (Sigma-Aldrich, St. Louis, MO, USA), complemented with the same supplements used in the previously described DMEM medium. All cells were kept at 37 °C and 5% CO2 in a Galaxy 170 R humidified incubator (Eppendorf, Hamburg, Germany). All cell lines have been routinely examined for mycoplasma contamination by using SPY-555-DNA (1:100 000, Spirochrome, Stein am Rhein, Switzerland) and by checking if any extracellular DNA is present. All cell lines used for experiments have been confirmed to be mycoplasma free.

### RNAi treatment

Cells for RNAi transfection were seeded on 35 mm uncoated dishes with the 0.17 mm (1.5 coverslip) glass thickness (Ibidi GmbH, Gräfeling, Germany) in 1 mL of the previously described DMEM medium to reach approximately 60% confluence the following day. siRNA constructs were diluted in OPTI-MEM medium (Life Technologies, Waltham, MA, US) and RNAi treatments were performed using Lipofectamine RNAiMAX Reagent (Life Technologies, Waltham, MA, US) according to the instructions provided by the manufacturer.

The siRNA constructs and their final concentrations were: 100 nM CENP-E siRNA (L-003252-000010; Dharmacon, Lafayette, CO, USA), 100 nM Spindly/CCDC99 siRNA (L-016970-00-0010, Dharmacon, Lafayette, CO, USA), 100 nM KIF4A siRNA (sc-60888; Santa Cruz Biotechnology, Dallas, TX, USA), 100 nM Kid/KIF22 siRNA (4392420; Ambion, Austin, TX, USA). Cells were incubated for four hours with the transfection mixture followed by medium replacement into the previously described DMEM medium. All experiments were performed 24 hours after siRNA transfection.

### Plasmid transfection

To transiently express CENP-A in an RPE1 cell line stably expressing tubulin-EYFP and H2B-mRFP, 5 µg of pHTN HaloTag-CENP-A plasmid was used for transfection performed on Nucleofector Kit R with the Nucleofector 2b Device (Lonza) following manufacturer’s instructions. The Nucleofactor Y-001 program was used for the transfection of the RPE1 cell line. After transfection, around 500 000 cells were seeded on 35 mm uncoated dishes with the 0.17 mm (1.5 coverslip) glass thickness (Ibidi GmbH, Gräfeling, Germany) in 1 mL of the previously described DMEM medium. Transfection was performed 48 hours before imaging.

### Lentivirus transduction and cell line generation

To produce lentiviruses, HEK293T cells were seeded on 10 cm Petri dishes to reach 80% confluence the next day. Plasmid constructs were diluted in OPTI-MEM medium (Life Technologies, Waltham, MA, US) and transfections were performed using Lipofectamine 3000 Transfection Reagent (Life Technologies, Waltham, MA, US) according to the manufacturer’s protocol. The plasmid constructs and their final masses were as follows: 10 µg of psPAX2, 5 µg of psMD2.G, 15 µg of pLH sgRNA Chr1-telo or 15 µg of pLH sgRNA Chr9-cen. The cell medium was replaced with the previously described DMEM medium 24 hours after transfection. Lentiviruses containing sgRNA for Chr1-telo or Chr9-cen were harvested 48 hours after transfection by filtering the cell medium through 45 µm filters (No. 99745, TPP, Switzerland). RPE1 cells stably expressing dCas9-GFP-3xFKBP were seeded in 6 well plates (120 000 cells/well, Thermo Fisher Scientific, Waltham, MA, USA) the day before lentivirus transduction. Lentiviruses were added to cells together with 6 µg/ml of polybrene (TR-1003-G, Merck Milipore, Darmstadt, Germany) and incubated for 24 hours.

### Treatments with inhibitors

The following Eg5 inhibitors were used on a “one cell per dish” principle: STLC (final concentration 40 µM, 164739, Sigma-Aldrich, St. Louis, MO, USA) was added approximately 30 s before landing to cause spindle shortening; FCPT (final concentration 40 µM, a gift from Timothy Mitchison) was added at the end of centripetal approaching to block the spindle elongation; monastrol (final concentration 100 µM in RPE1 cells, HY-101071A, MedChemExpress, Monmouth Junction, USA) was added 30 s before landing and then washed out with a pre-warmed DMEM medium when spindle stopped shortening. Nocodazole (final concentration 20 nM in RPE1 cells, HY-13520, MedChemExpress, Monmouth Junction, USA), Mps1 inhibitor AZ3146 (final concentration 3 µM, HY-14710, MedChemExpress, Monmouth Junction, USA), Mps1 inhibitor Cpd-5 (final concentration 62.5 nM, a gift from R. Medema via G. J. P. L. Kops lab; (Klaasen et al., 2022; Koch et al., 2016)) and actin polymerization inhibitor latrunculin A (final concentration 2 µM, 428021, Sigma-Aldrich, St. Louis, MO, USA) were added approximately 45 minutes prior to imaging and cells within the dish were subsequently imaged for 4 hours. CENP-E inhibitor GSK923295 (MedChemExpress, Monmouth Junction, USA, IC50 value 3.2 nM) was added at a final concentration of 80 nM 1-4 hours before imaging.

For live-cell imaging of monastrol-treated OVSAHO cells, 15 µM monastrol was chosen as it was sufficient to produce slower spindle elongation without inducing monopolar arrest in the majority of cells. In MDA-MB-231 cells, 30 and 50 µM monastrol produced a similar result. Only the cells that formed a bipolar spindle within the first ten minutes after NEBD were analyzed. For fixed-cell imaging of nocodazole-treated cancer cells, 10 ng/mL (∼33 nM) of nocodazole was added 2 hours prior to fixation.

### Live-cell dyes

To identify nuclear envelope breakdown in RPE1 cells stably expressing CENP-A-GFP and Centrin1-GFP, the SPY-555 or SPY-595 DNA probe (1: 100 000, Spirochrome, Stein am Rhein, Switzerland) was added to the cells 45 minutes to 2 hours prior to imaging. When microtubules were visualized during live-cell imaging in the same cell line, the SPY-650-tubulin probe (1:50 000, Spirochrome, Stein am Rhein, Switzerland) was added to the cells 45 minutes to 2 hours prior to imaging. For the live-cell imaging of OVSAHO, MDA-MB-231, and Caco-2 cell lines, the SPY-555 DNA probe (1: 50 000, Spirochrome, Stein am Rhein, Switzerland) was added to the cells along with the SiR-tubulin probe (50 nM, Spirochrome, Stein am Rhein, Switzerland; (Lukinavičius et al., 2014)) 45 minutes prior to imaging. The cells were subsequently imaged for 1-3 hours. For STED live-cell imaging, the SiR-tubulin probe (100 nM, Spirochrome, Stein am Rhein, Switzerland; (Lukinavičius et al., 2014)) was added to the cells 15 minutes prior to imaging and cells were subsequently imaged for 45 minutes. To avoid efflux of the dyes, a broad-spectrum efflux pump inhibitor verapamil (Spirochrome, Stein am Rhein, Switzerland) was added at a final concentration of 1 μM to cells along with tubulin and/or DNA probes. For the live-cell imaging of RPE1 cells stably expressing EYFP-α-tubulin and H2B-mRFP, cells were incubated for 30 minutes with Janelia Fluor 635 HaloTag ligand (a final concentration of 20 nM, Promega, Madison, WI, USA) 24 hours before imaging to visualize the transiently expressed CENP-A-HaloTag.

### Immunofluorescence

To determine the RNAi efficiency, RPE1 cells stably expressing CENP-A-GFP and Centrin1-GFP were grown on 35 mm uncoated dishes with the 0.17 mm (1.5 coverslip) glass thickness (Ibidi GmbH, Gräfeling, Germany) and fixed by 1 ml of ice-cold methanol for 1 minute at –20 °C, except for Spindly/CCD99 immunostaining where cells were pre-extracted with 0,2% Triton in PHEM buffer (60 mM PIPES, 10 mM EGTA, 25 mM HEPES, 4 mM MgCl2) for 1 minute and subsequently fixed with 4% pre-warmed to 37 °C for 10 minutes. Cells for CENP-E and ZW10 immunostainings were pre-extracted for 20 s with 0.5% Triton in PEM buffer (0.1M PIPES, 1mM EGTA, 1mM MgCl2) followed by fixation by 1 ml of ice-cold methanol for 1 minute. Following fixation, cells were washed three times for 5 minutes with 1 ml of PBS and permeabilized with 0.5% Triton-X-100 in PBS for 15 minutes at room temperature. To block unspecific binding, cells were incubated in 1 ml of blocking buffer (1% normal goat serum (NGS)) for 1 hour at 4 °C. Cells were then washed three times for 5 minutes with 1 ml of PBS and incubated with 250 µl of primary antibody solution overnight at 4 °C. The primary antibodies were used as follows: rabbit anti-CENP-E (1:100 for determination of RNAi efficiency, and 1:500 for staining of outer corona, C7488; Sigma-Aldrich), rabbit anti-Spindly/CCD99 (1:1000, A301_354A, Bethyl), mouse anti-KIF4A (1:100, sc-365144; Santa Cruz Biotechnology), mouse anti-Kid (1:100, sc-390640; Santa Cruz Biotechnology), rabbit polyclonal anti-ZW10 (1:500, ab21582, Abcam), rabbit polyclonal anti-MAD2 (1:250, 924601, BioLegend), mouse monoclonal anti-Astrin, clone C-1 (1:250, MABN2487, Merck). After the incubation with a primary antibody, cells were washed three times for 5 minutes with 1 ml of PBS and then incubated with 250 µl of secondary antibody for 1 hour at room temperature. Alexa Fluor 594 (Abcam, ab150108 or ab150076) was used as a secondary antibody at a 1:250 dilution for CENP-E, Kif4A, and Kid staining and at a 1:1000 dilution for Spindly staining in experiments where RNAi efficiency was tested. Alexa Fluor 647 (Abcam, ab150107) was used as a secondary antibody at a 1:500 dilution for Astrin staining. Alexa Fluor 594 was used as a secondary antibody at a 1:500 dilution for staining of CENP-E and Mad2 on polar kinetochores. Alexa Fluor 647 (Abcam, ab150075) was used as a secondary antibody at a 1:1000 dilution for ZW10 staining.

To image polar chromosomes in cancer cell lines OVSAHO, MDA-MB-231, and Caco-2, cells were fixed with 1 ml of 4% paraformaldehyde pre-warmed to 37 °C for 10 minutes followed by all the previously described steps for immunostaining. The primary antibodies were used as follows: rat anti-tubulin (1:500, MA1-80017, Invitrogen) and human anti-centromere protein (1:300, 15-234, Antibodies Incorporated, CA, USA). The secondary antibodies were: donkey anti-rat Alexa Fluor 488 (1:1000, Abcam, ab150153) and goat anti-human DyLight 594 (1:500, Abcam, ab96909). DAPI (1 µg/mL) was used for chromosome visualization in all cell lines.

For STED imaging, RPE1 cells stably expressing CENP-A-GFP and RPE1 cells stably expressing CENP-A-Cerulean and Mad2-mRuby were grown on 35 mm uncoated dishes with the 0.17 mm (1.5 coverslip) glass thickness (Ibidi GmbH, Gräfeling, Germany). Cell medium was removed and 0.5% Triton in PEM buffer (0.1M PIPES, 1mM EGTA, 1mM MgCl2) was added for 15 seconds to remove the components of the cytoplasm. Following extraction, cells were fixed in 3% paraformaldehyde and 0.1% glutaraldehyde solution for 10 minutes. To reduce the background fluorescence, quenching (100 mM glycine in PBS) and reduction (0.1% sodium borohydride in PBS) solutions were added for 7 and 10 minutes, respectively. To prevent non-specific binding, cells were incubated in blocking/permeabilization buffer (2% NGS and 0.5% Triton-X-100 in PBS) for 2 hours at 4 °C. Microtubules were then stained using the rat anti-tubulin primary antibody (1:500 in blocking/permeabilization buffer, MA1-80017, Invitrogen) with a 4 °C overnight incubation. The next day, cells were washed with PBS three times for 5 minutes. After washing, the secondary antibody donkey anti-rat Alexa Fluor 594 (1:1000 in blocking/permeabilization buffer, Abcam, ab150156) or donkey anti-rat Alexa Fluor 647 (1:1000 in blocking/permeabilization buffer, Abcam, ab150155) was added and incubated for 1 hour at room temperature. DAPI (1 µg/mL) or SPY-505-DNA (1:10 000, Spirochrome, Stein am Rhein, Switzerland) was added to visualize chromosomes. The protocol was originally modified for (Štimac et al., 2022) based on the protocols from (Ponjavić et al., 2021; Zhang et al., 2020). For validation of actin depolymerization by Latrunculin A, RPE1 cells stably expressing CENP-A-GFP and Centrin1-GFP were fixed following the same fixation protocol as above, and SiR-actin (200 nM in PBS, Spirochrome, Stein am Rhein, Switzerland) was incubated for 20 min at room temperature.

To analyze the position of chromosomes 1 and 9 in interphase cells, RPE1 cells stably expressing Chr1-telo-GFP or Chr9-cen-GFP were stained for tubulin and centrin-3 following the same immunostaining protocol used for STED imaging. Rabbit anti-centrin-3 primary antibody (1:1000, ab228690, Abcam) and secondary donkey anti-rabbit Alexa Fluor 647 (1:1000, Abcam, ab150075) were used for immunostaining.

### Microscopy

STED microscopy of fixed cells, live-cell imaging of pivoting microtubules, live-cell imaging of EB3 comets and live-cell imaging of cells treated with Cpd-5 were performed using an Expert Line easy3D STED microscope system (Abberior Instruments, Göttingen, Germany) with the 100 x/1.4NA UPLSAPO100x oil objective (Olympus, Tokio, Japan) and an avalanche photodiode (APD) detector, as previously described (Koprivec et al., 2025). The 405 nm, 488 nm, 561 nm and 647 nm laser lines were used for the excitation of Cerulean, GFP, Alexa Fluor 594/mRuby/mCherry and Alexa Fluor 647/Janelia Fluor 635/SiR-tubulin, respectively. The 775 nm laser line was used for the depletion of red and far-red lines during the STED superresolution imaging. Images were acquired using the Imspector software. The xy pixel size for fixed RPE1 cells stably expressing CENP-A-GFP was 20 nm and 20-30 focal planes were acquired with the 300 nm distance between them to cover the whole cell. The xy pixel size for fixed RPE1 cells stably expressing CENP-A-Cerulean and Mad2-mRuby was 25 nm and 1 focal plane containing the kinetochores of polar chromosomes was acquired. For live-cell STED imaging of pivoting microtubules, the xy pixel size was set to 30 nm and one focal plane was acquired every 60 seconds. For confocal live-cell imaging of pivoting microtubules in RPE1 cells on the same microscope, the xy pixel size was set to 55 nm and 3 focal planes were acquired, with the 1 µm distance between the planes and 15 seconds time intervals between different time frames. For confocal live-cell imaging of EB3 comets in RPE1 cells, the xy pixel size was set to 80 nm and 3 focal planes were acquired, with the 1 µm distance between the planes and 20 seconds time intervals between different time frames. For confocal live-cell imaging of EB3 comets in U2OS cells, the xy pixel size was set to 80 nm and one focal planes was acquired, with 1 second time intervals between different time frames. For confocal live-cell imaging of cells treated with Cpd-5, the 488 nm laser line was used to excite the GFP, the 647 nm laser line was used to excite the SPY-650-tubulin, the xy pixel size was 80 nm and 6-7 focal planes were acquired, with 1 µm distance between the planes and 20 seconds time intervals between different time frames, as previously described (Klaasen et al., 2022). Movies of U2OS cells expressing CENP-A-GFP and tubulin-mCherry were acquired as previously described (Klaasen et al., 2022).

Confocal microscopy of OVSAHO cells and RPE1 cells immunostained to check the respective RNAi efficiencies was performed using a previously described microscope setup (Buđa et al., 2017), consisting of a Bruker Opterra Multipoint Scanning Confocal Microscope (Bruker Nano Surfaces, Middleton, WI, USA), mounted on a Nikon Ti-E inverted microscope with a Nikon CFI Plan Apo VC 100 x/1.4 numerical aperture oil objective (Nikon, Tokyo, Japan). To excite DAPI, GFP, Alexa Fluor 594 or DyLight 594, a 405 nm, 488 nm and 561 nm laser lines were used, respectively. Opterra Dichroic and Barrier Filter Set 405/488/561 enabled the separation of excitation light from the emitted fluorescence. Images were acquired using the Evolve 512 Delta Electron Multiplying Charge Coupled Device (EMCCD) Camera (Photometrics, Tuscon, AZ, USA), with a camera readout mode of 20 MHz. The xy pixel size was 83 nm and the slit aperture was 22 µm. In all experiments, the whole spindle stack was imaged. The z-stacks were acquired using the unidirectional xyz scan mode with the 1 μm distance between the planes to cover the entire cell.

Confocal live cell imaging of RPE1 cells stably expressing CENP-A-GFP and Centrin1-GFP treated with STLC, FCPT, monastrol and AZ3146 was performed on a Dragonfly spinning disk confocal microscope system (Andor Technology, Belfast, UK) using the 100x/1.47 HC PL APO oil objective (Nikon, Tokyo, Japan) and Sona 4.2B-6 Back-illuminated sCMOS camera (acquisition mode set to high speed, Andor Technology) linked with iXon Life 888 EMCCD Camera (EM gain set to 208, Andor Technology). To simultaneously image two channels on two independent cameras, the 565 LP image splitter was used. Images were acquired using the Fusion software. During imaging, cells were maintained at 37°C and 5% CO2 within the heating chamber (Okolab, Pozzuoli, NA, Italy). For the excitation of GFP and SPY-650-tubulin, the 488-nm and 637-nm laser lines with corresponding filters were used, respectively. Image acquisition was done every 0.4 µm with 21 focal planes and a time interval of 10 seconds.

The Lattice Lightsheet 7 system (Carl Zeiss, Germany) was used for the live cell imaging of control RPE1 cells expressing CENP-A-GFP and Centrin1-GFP and for the imaging of cells with CENP-E, Spindly/CCD99 and Kid/KIF4A depletion, inhibition of actin polymerization or nocodazole addition. The same system was used for live-cell imaging of cancer cell lines. The system includes an illumination objective lens 13.3×/0.4 (at a 30° angle to cover the glass) with a static phase element and a detection objective lens 44.83×/1.0 (at a 60° angle to cover the glass) with an Alvarez manipulator, as previously described (Vukušić and Tolić, 2024). For the live-cell imaging of RPE1 cells, the GFP was excited using the 488-nm diode laser (power output 10 Mw), with the laser power set to 2.5%, LBF 405/488/561/642 emission filter and 12 ms exposure time. The detection module consisted of a Hamamatsu ORCA-Fusion sCMOS camera. The width of the imaging area was set to 32-36 µm with the x interval of 0.3 µm. Images were acquired every 6 seconds. For the live-cell imaging of cancer cell lines, the SPY-555-DNA was excited using the 561-nm diode laser, with the laser power set to 2.0-4.0%. SiR-tubulin was excited using the 561-nm diode laser, with the laser power set to 1.0-1.5%. The exposure time was 15-20 ms. The width of the imaging area was set to 36 µm with the x interval of 0.5 µm. Images were acquired every minute. During the imaging, cells were kept at 37 °C and at 5% CO2 in a Zeiss stage incubation chamber system (Carl Zeiss, Germany). All movies were deskewed with the ZEN 3.7 software using the “Linear Interpolation” and „Cover Glass Transformation” settings. Deconvolution was performed using the „Constrained Iterative” algorithm with automatic normalization and strength set to 4.0. The imaging parameters for CENP-E inhibition experiments were described previously (Vukušić and Tolić, 2024).

Confocal imaging of chromosomes 1 and 9 in RPE1 cells was performed on the Airyscan Zeiss LSM800 confocal laser scanning microscope with the 40x/0.95 Corr M27 objective (Carl Zeiss, Germany) and LSM 800 camera. To excite DAPI, GFP, Alexa Fluor 594 and 647, the 405 nm, 488 nm, 561 nm and 647 nm laser lines were used, respectively. Images were acquired in ZEN blue 3.5 software using the Tiles recording mode, with the 0.5 µm distance between the planes to cover the whole cells.

### Image analysis

In lattice light-sheet and confocal videos, kinetochores as markers for chromosomes and centrosomes as markers for spindle poles were tracked in a z-stack using the Point Tool in Fiji/ImageJ. When the signal covered several z-planes, the central z-plane was used for tracking. Spindle length, pivoting angle (i.e., the angle between the long spindle axis and the line that connects the kinetochore midpoint with the closer centrosome), cTilt (i.e., the angle between the long spindle axis and the line that connects sister kinetochores; (Renda et al., 2022)), interkinetochore distance, distance to closer and further centrosome and distance to centrosome midpoint were subsequently calculated from the coordinates in 3D using a home-written MatLab script. The end of centripetal movement was determined as a time point when the distance of the kinetochore midpoint to the closer centrosome started varying instead of continuously becoming smaller. The landing on the main spindle surface was determined as a time point when the pivoting angle was below 60° and cTilt was below 22.5° (Renda et al., 2022), corresponding to the moment when kinetochores first align along the microtubules of the main spindle surface, even if they experience subsequent detachments. The landing was additionally verified visually by checking that a kinetochore pair was adjacent to the microtubules of the main spindle body. The alignment was defined as the moment when the kinetochore pair was equidistant (±1-2 µm) to both poles. The formation of the prometaphase ring was visually determined as a time point when most kinetochores finish their centripetal movement. The end of spindle elongation was defined as a time point when spindle length starts varying and/or decreasing instead of continuously increasing. The anaphase onset was visually determined as a time point when sister kinetochores first start separating. Polar chromosomes were defined as those that had a pivoting angle of >90° at NEBD and subsequently made their first contact with the spindle behind the spindle pole. The position of the first contact was determined at the end of centripetal movement, when the distance of the kinetochore pair to the spindle contour was usually <3 µm. Pivoting was then analyzed between the end of centripetal movement until after landing. Only kinetochores that could be reliably tracked from NEBD to alignment, remaining distinguishable from neighboring kinetochores throughout this period, were included in the analysis. In Eg5 manipulation experiments, analysis was done from the first frame when the inhibitor started working, i.e., when the spindle length started increasing or decreasing.

Successful passage across the pole was defined as the passage across the 90° mark within 15 minutes from NEBD, not taking into account any subsequent failures to form stable attachments to microtubules of the main spindle surface. Misaligned chromosomes were defined as those located within the two spindle poles and less than 2 µm away from the centrosome/spindle pole in metaphase. Unaligned chromosomes were defined as those located behind the spindle pole or, alternatively, in front of the pole but not attached to microtubules of the main spindle surface in metaphase. Lagging chromosomes were defined as those located in the central part of the spindle with a stretched kinetochore during anaphase.

When chromosomes were labeled instead of kinetochores, two points were put in the centromeric region during the tracking. When poles were marked with tubulin or EB3 instead of Centrin1, the point was put in the center of tubulin/EB3 signal. When astral microtubules were tracked instead of chromosomes or kinetochores, their movement was tracked by putting two points along the central part of the microtubule contour and their shape was determined by putting two points at the spindle poles and subsequent five along the entire microtubule contour. Kinetochore switching between microtubules was investigated using visual analysis.

In STED images of fixed cells, NEBD was determined as the stage when the centrosomes were one above the other, or when the centrosomes were positioned sideways, but microtubules did not occupy the whole area between the centrosomes. The beginning of the centripetal movement was identified by the apparent attachment of all kinetochores to microtubules at varying distances from the centrosomes, including distances larger than 4 µm. The end of the centripetal movement was determined as the stage when kinetochores were arranged in a prometaphase ring at similar distances, smaller than 4 µm, to the centrosomes. Passage across the pole was identified by the attachment of a kinetochore pair to the microtubules that were not part of the main spindle body at a stage when the spindle was almost fully formed. Landing was determined as the stage when a kinetochore pair was found adjacent to the microtubules of the main spindle body less than 4 µm from the centrosome, and traveling to the equator when the kinetochores were between 4 µm away from the centrosome and 2 µm away from the equator.

The length of long xy-, short xy- and z nuclear axes was measured on maximum intensity projections of the whole cell just before NEBD using the Line Tool in ImageJ. The long xy axis was defined as the axis with the largest distance between the two opposite sides of the nuclear border, the short xy axis was defined as perpendicular to the middle of the long xy axis and z axis was defined as the distance from the first focal plane in which the signal was present until the last. The spindle axis, used to determine the danger zone, was measured using the Line Tool in ImageJ on maximum intensity projections of the whole cell when the prometaphase ring formed and most kinetochores finished their centripetal movement. The angle between the long nuclear xy axis just before NEBD and the long spindle axis in prometaphase was then calculated to determine which chromosomes are most likely to attach behind the pole. This angle was also compared to the angle between the long cellular xy axis at NEBD, determined using the Line Tool in ImageJ, and the long spindle axis in prometaphase.

The long axis of interphase RPE1 dCas9-GFP cells with labeled subtelomeric region of chromosome 1 or pericentromeric region of chromosome 9 was measured on maximum intensity projections of the whole cell. The long axis was then divided into three equal parts that represent the borders of three cellular compartments - two polar zones and the central zone. The position of the labeled chromosomes was determined as polar if they were located in any of the polar zones and as central if they were located in the central zone, respectively.

To determine the position of kinetochore pairs relative to the pole in STED images, kinetochores and poles were marked in 3D using a Point Tool in ImageJ and the angle that the line connecting the kinetochore pair midpoint with the closer centrosome forms with the long spindle axis (i.e., pivoting angle) was subsequently calculated using a home-written MatLab script. Kinetochore pairs in fixed-cell STED images were considered polar if they were located above 90° or, alternatively, were located within ∼3 μm from the pole but not attached to the microtubules of the main spindle surface during later stages of prometaphase when the spindle was already formed.

Attachment types were determined from STED images of fixed cells. Polar chromosomes were considered attached to microtubules when there was a microtubule either directly ending at the kinetochore or in very close proximity to the kinetochore. Polar chromosomes were considered unattached when there was no microtubule in their proximity. The type of attachment was determined visually. A limitation of this method is that attachment cannot always be distinguished from mere proximity.

For fixed-cell confocal images of CENP-A together with Mad2 or Astrin or CENP-E or ZW10, protein levels were measured using the Polygon Tool in ImageJ. The same polygon was used for measuring the signal in both channels for the same kinetochore, as well as for the background in the vicinity of the kinetochore in both channels. The final Mad2/CENP-A signal on the kinetochore was calculated using the formula: Mad2/CENP-A signal = (Integrated density of Mad2 on the kinetochore – (Kinetochore area x Mean fluorescence of the background))/(Integrated density of CENP-A on the kinetochore – (Kinetochore area x Mean fluorescence of the background)). The other 3 proteins were analyzed in the same manner. In fixed-cell confocal images, kinetochore pairs were classified as polar if their angle exceeded 90°. For each cell, three to ten kinetochores located between the centrosomes were analyzed and classified as non-polar. Additionally, three to ten kinetochores per metaphase cell were analyzed. Metaphase cells were identified by the presence of fully formed metaphase plates.

The intensity of microtubules from the opposite side of the spindle, the intensity of astral microtubules and the accompanying background were measured in a single z-plane using the 20x20 pixel Square Tool in ImageJ. The final intensity of the microtubules was then calculated using the formula: Microtubule intensity = Mean intensity of the microtubule – Mean intensity of the background.

In the validation experiments for protein depletion by siRNA, CENP-E, Spindly, KIF4A and Kid intensities were measured in sum intensity projections of the entire cell, covering 33 z-planes or 33 µm. The total signal of the protein of interest within the spindle was marked by using the Polygon Tool in ImageJ. Background in the cytoplasm was measured using the 25x25 pixel Square Tool. The total intensities were calculated using the following formula: Protein intensity = Integrated density within the spindle – (Area of the spindle x Mean fluorescence of the background). The total intensities were then normalized to the mean total intensity obtained from control cells.

Tracking of kinetochores in U2OS cells expressing CENP-A-GFP and tubulin-mCherry was performed using 2D maximum intensity projections and the ‘TrackMate’ plugin in ImageJ, with the LoG detector (Ershov et al., 2022). The estimated blob diameter was set to match the size of a single kinetochore labeled with CENP-A, and thresholding was adjusted until all visible kinetochores were detected, with no false positives outside the cell. To avoid interference from neighboring cells, the field of view was cropped accordingly. Using this approach, the average number of tracked kinetochore pairs at nuclear envelope breakdown (NEBD) was 42, fewer than the reported modal chromosome number for the U2OS cell line (n = 74, ATCC). This discrepancy is likely due to insufficient spatial resolution in 2D projections, which may prevent the complete separation of overlapping kinetochore pairs. The number of polar chromosomes that successfully passed the spindle pole and reached the metaphase plate by the end of spindle elongation in live-imaged U2OS cells was calculated by subtracting the number of identified polar kinetochores remaining behind the pole at the end of spindle elongation from the total number of polar kinetochores counted at NEBD. Kinetochore pairs at NEBD and at the end of spindle elongation were classified as polar if their pivoting angle in 2D projection exceeded 90°. The end of spindle elongation was estimated as described previously (Klaasen et al., 2022). The majority of kinetochores remaining behind the pole were continuously backtracked frame by frame to NEBD, enabling clear distinction from those that successfully traversed the spindle pole.

For live-cell measurements of spindle elongation and polar chromosome transit in cancer cell lines, the spindle length was measured each minute as the 2D distance between the two spindle poles using the Line Tool in ImageJ. Coupled with this were the measurements of the polar transit time for each cell, defined as the time point when the last chromosome of those that were initially polar moved from the back side to the front side of the centrosome.

All measurements were performed in ImageJ/Fiji (National Institutes of Health), followed by data analysis in MATLAB (MathWorks). Figures and schematic representations were assembled in Adobe Illustrator CC (Adobe Systems). Generally, the Shapiro-Wilk test was used to determine normality of the sample data. T-test (for two samples unless stated otherwise) or ANOVA with post-hoc Tukey test (for more than two samples) were then used for data that followed a normal distribution, and Mann-Whitney U test (for two samples) or Kruskal-Wallis with posthoc Dunn’s test (for more than two samples) were used otherwise. To analyze categorical data, Chi-squared test was used for larger samples (N > 50) and Fisher’s exact test was used for smaller samples. The result was considered statistically significant when the p-value was equal to or less than 0.05. Statistical analysis was performed as noted in the respective Figure captions.

## Author Contribution

I.K. and V.Š. performed all experiments and analyzed the data, except for the following: experiments on Astrin and the cancer cell lines MDA-MB-231, Caco-2, and a subset of OVSAHO cells were conducted by M.Đ., experiments involving CENP-E inhibition, corona proteins, and U2OS cells with labeled kinetochores and microtubules were carried out by K.V., and experiments on chromosomes 1 and 9 were performed by P.M. I.K., V.Š., and I.M.T. wrote the manuscript with contribution from M.Đ. and K.V. I.M.T. supervised the project.

## Acknowledgments

The authors thank Alexey Khodjakov, Jonathon Pines, Andrew Holland, Will Krek, Marin Barišić, Susanne Lens and Julie Welburn for the cell lines; Oliver Vugrek and Filip Rokić for help with lentiviral work, Marko Šprem and the Abberior team for help with the Expert Line easy3D STED microscope system setup and developing microscopy protocols; Ivana Šarić for help with the drawings, members of the Tolić, Pavin, Kops, and Meraldi groups for helpful discussions and advice. The authors acknowledge funding by the European Research Council (ERC Synergy Grant, GA Number 855158), the Croatian Science Foundation (HRZZ) through Swiss-Croatian Bilateral Projects (project IPCH-2022-10-9344) and Cooperation Programme with Croatian Scientists in the Diaspora "Research Cooperability" (project PZS-2019-02-7653), and projects co-financed by the Croatian Government and the European Union through the European Regional Development Fund—the Competitiveness and Cohesion Operational Programme: IPSted (Grant KK.01.1.1.04.0057) and QuantiXLie Center of Excellence (Grant KK.01.1.1.01.0004).

## Competing interests

The authors declare no competing interests.

## Data availability

Source data are provided with this paper. All other data supporting the findings of this study are available from the corresponding author on reasonable request.

## Code availability

Codes used to analyze and plot the data are available from the corresponding author on request.

## Supplementary Material

### Supplementary Figures S1-S7

**Supplementary Figure 1.**
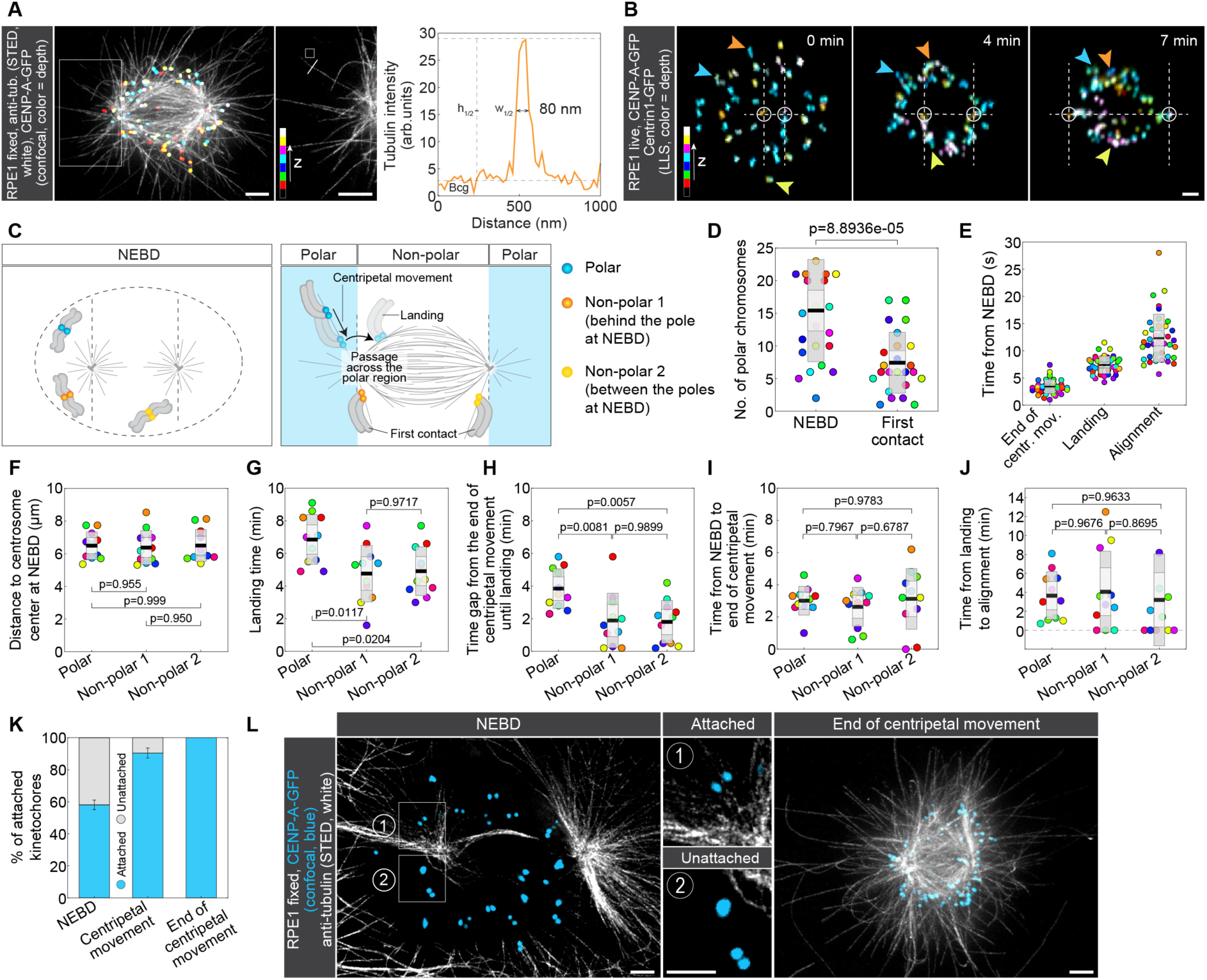
**(A)** STED superresolution images of microtubules immunostained for α-tubulin (white) in RPE1 cells stably expressing CENP-A-GFP (colorful, confocal). Images show maximum intensity projection of the whole cell (left) and the inset of a single z-plane with the astral microtubule of interest (right), marked with the white square on the left image. Kinetochores are color-coded for depth with the brgbcmyw LUT in ImageJ throughout 12 z-planes, correspoding to 3.6 µm. Line Tool (length = 1 µm, thickness = 1) and Square Tool (25x25 pixel, corresponding to 0.5x0.5 µm) from ImageJ are drawn on the inset and represent tools to measure the intensity profile of the astral microtubule and the mean intensity of the nearby background, respectively. The graph on the right shows the intensity profile of the line drawn perpendicularly to the astral microtubule in the STED image inset. The resolution is defined as the width of the peak at its half-maximum, after subtracting the background. **(B)** Time-lapse LLS images of RPE1 cells stably expressing CENP-A-GFP and Centrin1-GFP (colorful, lattice light-sheet). Images show maximum intensity projections of 29, 34 and 40 z-planes (4.2-5.8 µm) covering the whole cell, respectively. The blue arrow represents a polar kinetochore, orange and yellow arrows represent non-polar kinetochores located behind the pole or between the poles at NEBD, respectively. All kinetochores are tracked from NEBD until alignment to the spindle equator. Kinetochores are color-coded for depth with the brgbcmyw LUT in ImageJ. **(C)** Schematic representation of three groups of kinetochores. Blue kinetochores represent polar kinetochores located behind the spindle pole at NEBD and at their first contact with the spindle. Orange kinetochores represent non-polar kinetochores that are located behind the spindle pole at NEBD but between the two poles when they make their first contact with the spindle. Yellow kinetochores represent non-polar kinetochores located between the spindle poles at NEBD and during their first contact with the spindle. Colors for each group correspond to arrows in (B). **(D)** Univariate scatter plot of the number of chromosomes located behind the pole at NEBD and at their first contact with the spindle. Each color represents the same cell at each time point. N = 24 cells from 8 experiments. Statistical analysis: t-test. **(E)** Univariate scatter plot of the time from NEBD until the end of centripetal movement, landing and alignment. Each color represents the same kinetochore pair. N = 39 kinetochore pairs in 20 cells from 8 experiments. **(F-J)** Univariate scatter plot of the (F) distance to centrosome center at NEBD, (G) landing time, (H) time gap from the end of centripetal movement until landing, (I) time from NEBD to centripetal movement, and (J) time from landing to alignment for the three groups of kinetochores described in (C). Each color represents one kinetochore pair. N = 11 kinetochore pairs in 10 cells from 7 experiments. Statistical tests: Anova with post-hoc Tukey test. **(K)** The fractions of kinetochore pairs next to microtubules (blue) and without microtubules in their proximity (gray) during NEBD, centripetal movement and at the end of centripetal movement. N = 241 kinetochore pairs in 15 cells from 6 experiments (NEBD), 73 kinetochore pairs in 4 cells from 3 experiments (centripetal movement), 11 cells from 4 experiments, where all kinetochore pairs located behind the pole had a microtubule attached to them (end of centripetal approaching). **(L)** STED superresolution images of microtubules immunostained for α-tubulin (white) in RPE1 cells stably expressing CENP-A-GFP (blue, confocal) at NEBD and at the end of centripetal movement. Images show a single z-plane at NEBD and a maximum intensity projection of 23 z-planes (6.9 µm) covering the whole cell at the end of centripetal movement. Two types of attachments (attached vs. unattached) are shown in insets from NEBD. **(D-J)** Boxes represent standard deviation (dark gray), 95% confidence interval of the mean (light gray) and mean value (black). **(A,B,L)** Scale bars, 2 µm.

**Supplementary Figure 2.**
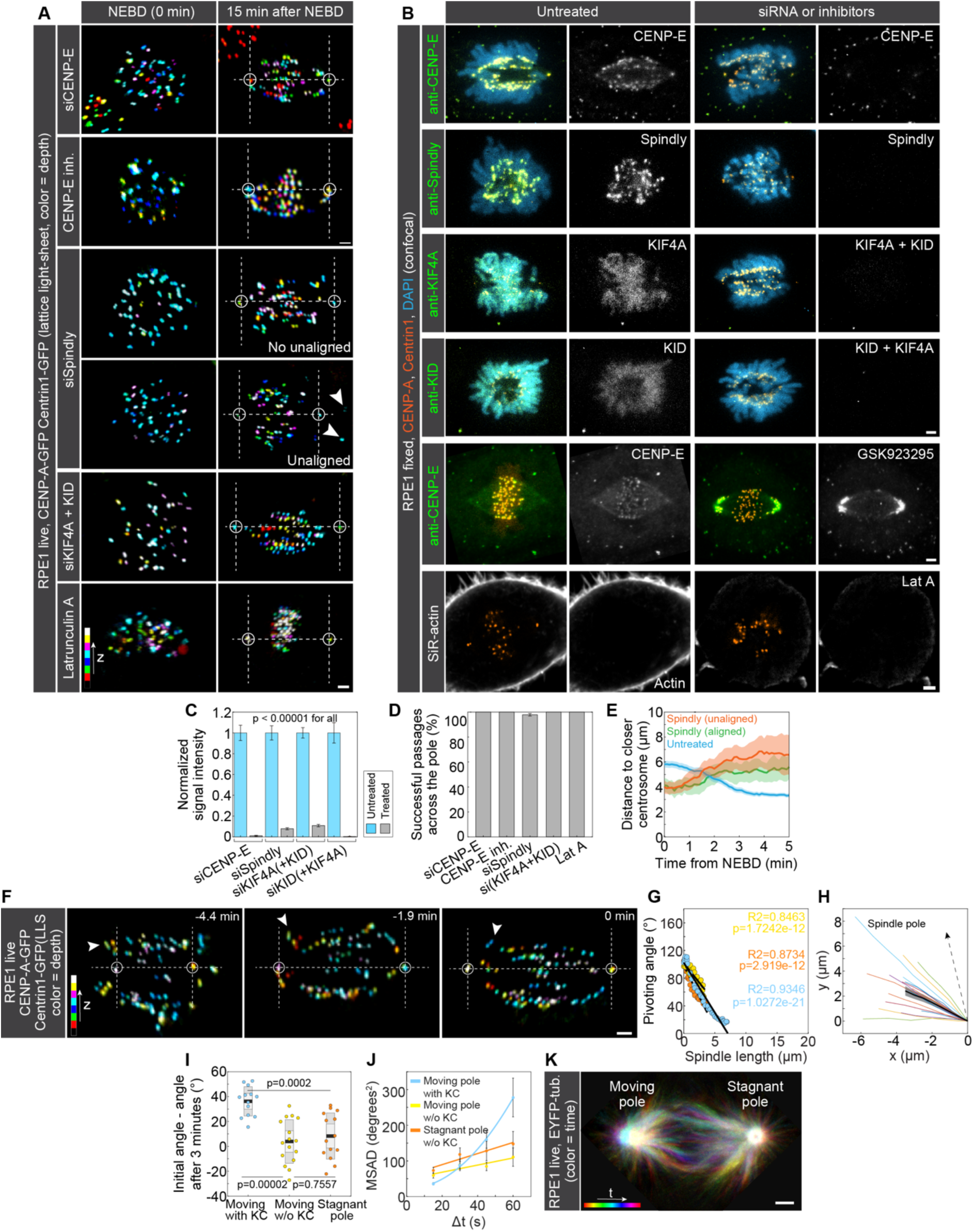
**(A)** Time-lapse images of RPE1 cells stably expressing CENP-A-GFP and Centrin1-GFP (colorful, lattice light-sheet) after CENP-E depletion by siRNA, CENP-E inhibition by 80 nM GSK923295, Spindly depletion by siRNA, KIF4A/Kid co-depletion by siRNA, and actin depolymerization by Latrunculin A treatment. Images show maximum intensity projections of the whole cell. Kinetochores are color-coded for depth with the brgbcmyw LUT in ImageJ throughout 27-77 z-planes, corresponding to 3.9-11.2 µm. White arrowheads mark unaligned kinetochores after Spindly depletion. **(B)** Validation of protein depletion by siRNA, CENP-E inhibition, and actin depolymerization. Columns 1 and 2 show untreated cells, and columns 3 and 4 treated cells. Columns 1 and 3 display merged channels of confocal images, and columns 2 and 4 only the protein of interest. Cells were immunostained for the proteins of interest denoted in images in column 2 (shown in green in merged images, except for actin which was stained by SiR-actin and shown in white), and the treatments are denoted in images in column 4. RPE1 cells stably expressing CENP-A-GFP and Centrin1-GFP (both in orange) were used, with chromosomes stained with DAPI (blue, rows 1-4), except in row 5, where RPE1 cells stably expressing CENP-A-GFP were used. Images are maximum intensity projections of the whole cell (33 z-planes), corresponding to 16.5 µm, except row 6, which is a single plane. Note that inhibition of actin polymerization by latrunculin A was additionally validated by retraction and rounding of interphase cells. **(C)** Normalized signal intensity for siRNA treatments in (B) for untreated (blue) and treated (gray) cells. N = 36 control and 36 CENP-E siRNA-treated cells from 3 experiments. N = 32 control and 30 Spindly siRNA-treated cells from 3 experiments. N = 37 control and 35 KIF4A/Kid siRNA-treated cells from 3 experiments stained with KIF4A antibody. N = 33 control and 30 KIF4A/Kid siRNA-treated cells from 3 experiments stained with Kid antibody. Statistical test: Mann-Whitney U test. **(D)** Fraction of successful passages of polar kinetochore pairs across the spindle pole in cells treated with CENP-E siRNA, CENP-E inhibitor GSK923295, Spindly siRNA, KIF4A+Kid siRNA, and latrunculin A. Successful passage was defined as passage across the 90° mark with respect to the spindle axis within 15 minutes from NEBD. All polar kinetochores moved across the pole within 15 minutes from NEBD in all treatments, except for Spindly depletion. N >150 polar kinetochore pairs in 11 cells from 3 experiments for CENP-E siRNA. N >300 polar kinetochore pairs in 22 cells from 2 experiments for CENP-E inhibition by GSK923295. N = 188 polar kinetochore pairs in 12 cells from 6 experiments for Spindly siRNA, 4 out of which failed to pass the pole. N >100 polar kinetochore pairs in 10 cells from 3 experiments for KIF4A+Kid siRNA. N >100 polar kinetochore pairs in 10 cells from 4 experiments for Latrunculin A. Note that, for cases in which all polar kinetochores crossed the pole within 15 minutes of NEBD, we did not directly count kinetochores per cell, but instead estimated a conservative lower bound based on the average number of polar kinetochores per cell at NEBD. **(E)** Distance of a polar kinetochore pair to the closer centrosome in time for polar kinetochore pairs in untreated cells (blue), polar kinetochore pairs in Spindly depletion that successfully crossed the pole (green) and polar kinetochore pairs that failed to cross the pole (orange). Values are shown as mean (dark line) and SEM (shaded areas). N = 39 polar kinetochore pairs in 20 untreated cells from 8 experiments. N = 4 polar kinetochore pairs that successfully crossed the pole in 3 Spindly-depleted cells from 2 experiments. N = 4 polar kinetochore pairs that failed to cross the pole in 3 Spindly-depleted cells from 2 experiments. **(F)** Time-lapse images of RPE1 cells stably expressing CENP-A-GFP and Centrin1-GFP (colorful, lattice light-sheet). Images show maximum intensity projections of the whole cell. Kinetochores are color-coded for depth with the brgbcmyw LUT in ImageJ throughout 34-38 z-planes, corresponding to 4.9-5.5 µm. White arrows represent one polar kinetochore tracked from the end of centripetal movement until it passed the spindle pole and landed on the main spindle surface (t = 0 minutes). **(G)** The anticorrelation between the pivoting angle and the spindle length for three individual polar kinetochore pairs as in (F). Linear regression. N = 3 randomly chosen polar kinetochore pairs in 3 cells from 3 experiments. **(H)** The shape of astral microtubules with attached polar kinetochores tracked mid-pivoting. The spindle pole is located in (0,0). N = 16 microtubules in 10 cells from 4 experiments. Values are shown as mean (dark line) and SEM (shaded areas) and individual microtubules are depicted in colors. **(I)** Pivoting angle change over a period of 3 minutes from the beginning of pivoting for astral microtubules with polar kinetochores attached to them on the moving pole (blue), microtubules without polar kinetochores attached to them on the moving pole (yellow) and microtubules without polar kinetochores attached to them on the stagnant pole (orange). N = 13 microtubules with polar kinetochores attached to them on the moving pole in 9 cells from 4 experiments. N = 16 microtubules without polar kinetochores attached to them on the moving pole in 10 cells from 4 experiments. N = 14 microtubules without polar kinetochores attached to them on the stagnant pole in 6 cells from 4 experiments. Boxes represent standard deviation (dark gray), 95% confidence interval of the mean (light gray) and mean value (black). Statistical test: Anova with post-hoc Tukey test. **(J)** Mean squared angular displacement (MSAD) for astral microtubules with polar kinetochores attached to them on the moving pole (blue), microtubules without polar kinetochores attached to them on the moving pole (yellow) and microtubules without polar kinetochores attached to them on the stagnant pole (orange). MSAD = v^2^t^2^ + 2D_MT_Δt + offset for microtubules with polar kinetochores attached to them on the moving pole and MSAD = 2D_MT_Δt + offset for the other two groups of microtubule (Berg, 1993). R2 = 0.999, 0.841 and 0.913, respectively. Error bars represent SEM. N = 16 microtubules with polar kinetochores attached to them on the moving pole in 10 cells from 4 experiments. N = 16 microtubules without polar kinetochores attached to them on the moving pole in 10 cells from 4 experiments. N = 14 microtubules without polar kinetochores attached to them on the stagnant pole in 6 cells from 4 experiments. **(K)** An image of an RPE1 cell stably expressing EYFP-tubulin (colorful, confocal) and H2B-mRFP (not imaged) with added HaloTag-CENP-A (not shown). The image is a temporal maximum intensity projection of the spindle and centrosome movements over 3 central z-planes (3 µm) and a time period of 4.5 minutes, color-coded for time with the Spectrum LUT in ImageJ. **(A,B,F,K)** Scale bars, 2 µm.

**Supplementary Figure 3.**
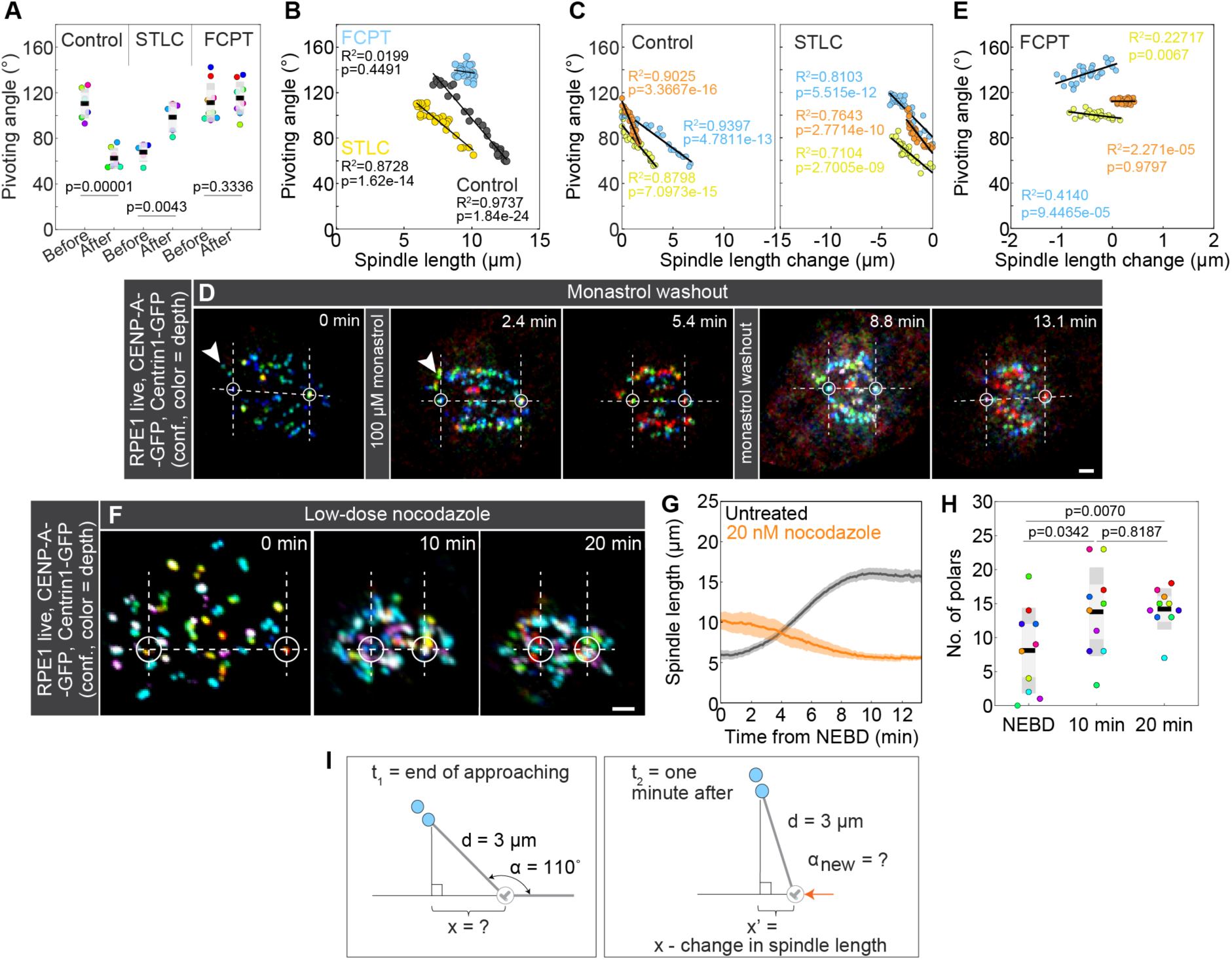
**(A)** Univariate scatter plot of the pivoting angle before and 5 minutes after the addition of DMSO (left), STLC (middle) and FCPT (right). Each color represents a different kinetochore pair. If landing occurred less than 5 minutes after the addition of DMSO, it was taken as the final step instead. N = 7 kinetochore pairs in 7 cells from 7 experiments (DMSO), 5 kinetochore pairs in 3 cells from 3 experiments (STLC), 9 kinetochore pairs in 7 cells from 7 experiments (FCPT). Statistical test: paired t-test. **(B)** Anticorrelation of the pivoting angle and spindle length for the control, STLC and FCPT treatment. N = 1 kinetochore pair from 1 cell and 1 experiments for each. **(C)** Anticorrelations of the pivoting angle and spindle length change for three other kinetochore pairs in control (left) and STLC treatment. N = 3 kinetochore pairs in 3 cells from 3 experiments. **(D)** Time-lapse images of an RPE1 cell stably expressing CENP-A-GFP and Centrin1-GFP (colorful, confocal) for monastrol treatment and washout. White circles represent spindle poles, horizontal dashed lines represent the long spindle axis and lines perpendicular to it denote the borders of the polar region. Monastrol treatment caused spindle shortening and was reversed by washout. Kinetochores are color-coded for depth with the brgbcmyw LUT in ImageJ. **(E)** Pivoting angle versus spindle length change for three kinetochore pairs in FCPT treatment. N = 3 kinetochore pairs in 3 cells from 3 experiments. **(F)** Time-lapse lattice light-sheet images of RPE1 cells stably expressing CENP-A-GFP and Centrin1-GFP (colorful, confocal) in low-dose nocodazole treatment (20 nM). Images are maximum intensity projections of the whole cell. Kinetochores are color-coded for depth with the brgbcmyw LUT in ImageJ throughout 37-52 z-planes, corresponding to 5.4-7.5 µm. White circles represent spindle poles, dashed lines represent the long spindle axis and lines perpendicular to it denote the borders of the polar region. **(G)** Change of spindle length in time from NEBD for untreated (gray) and cells treated with 20 nM nocodazole (orange). Values are shown as mean (dark line) and SEM (shaded areas). N = 20 untreated cells from 8 experiments. N = 10 nocodazole-treated cells from 4 experiments. **(H)** The number of chromosomes behind the pole at NEBD and 10 and 20 minutes after NEBD for cells treated with 20 nM nocodazole. Each color represents one cell. N = 10 cells from 4 experiments. Statistical test: repeated measures Anova. **(I)** Schematic representation for calculating the theoretical line shown in Figure 3H. d = distance between the closer spindle pole and the kinetochore pair (measured in Supplementary Figure 2E), x = distance between the spindle pole and the orthogonal projection of the kinetochore pair onto the spindle axis, x’ = x – change in spindle length, α = pivoting angle, α_new_ = calculated as the arccos (x’/d). The values of d and α were obtained from the DMSO and STLC experiments. **(A,H)** Boxes represent the standard deviation (dark gray), 95% confidence interval of the mean (light gray) and mean value (black). **(B,C,E)** Linear regression. **(D,F)** Scale bars, 2 µm.

**Supplementary Figure 4.**
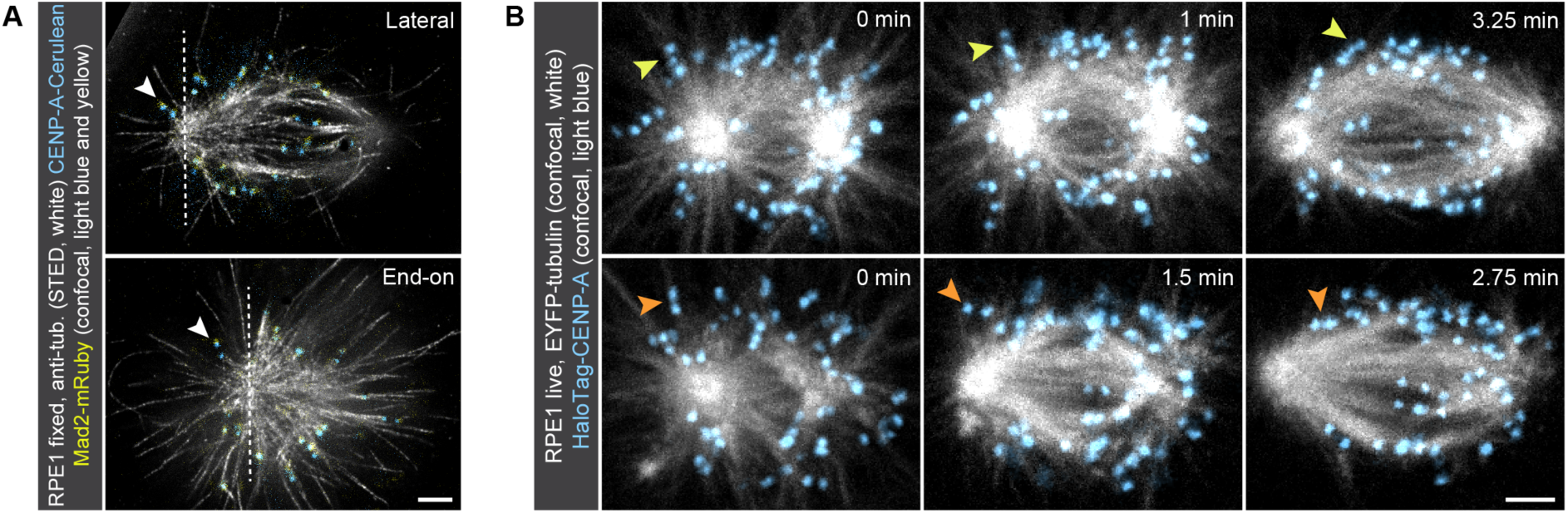
**(A)** STED superresolution images of microtubules immunostained for α-tubulin (white) in RPE1 cells stably expressing CENP-A-Cerulean (blue) and Mad2-mRuby (yellow). Images show single z-planes of spindles with laterally or end-on attached polar kinetochores shown in Figure 4. **(B)** Time-lapse confocal images of RPE1 cells stably expressing EYFP-α-tubulin (white) and H2B-mRFP (not imaged) with added HaloTag-CENP-A (blue). The first row represents a kinetochore pair that constantly remains attached to the same microtubule during pivoting. The second row represents a kinetochore pair that changes the attachment of its further kinetochore during pivoting from lateral to bilateral. The images are maximum intensity projections of 3 central z-planes spanning 3 µm. Kinetochores are smoothed with 0.75-mm-sigma Gaussian blur. **(A-B)** Scale bars, 2 µm.

**Supplementary Figure 5.**
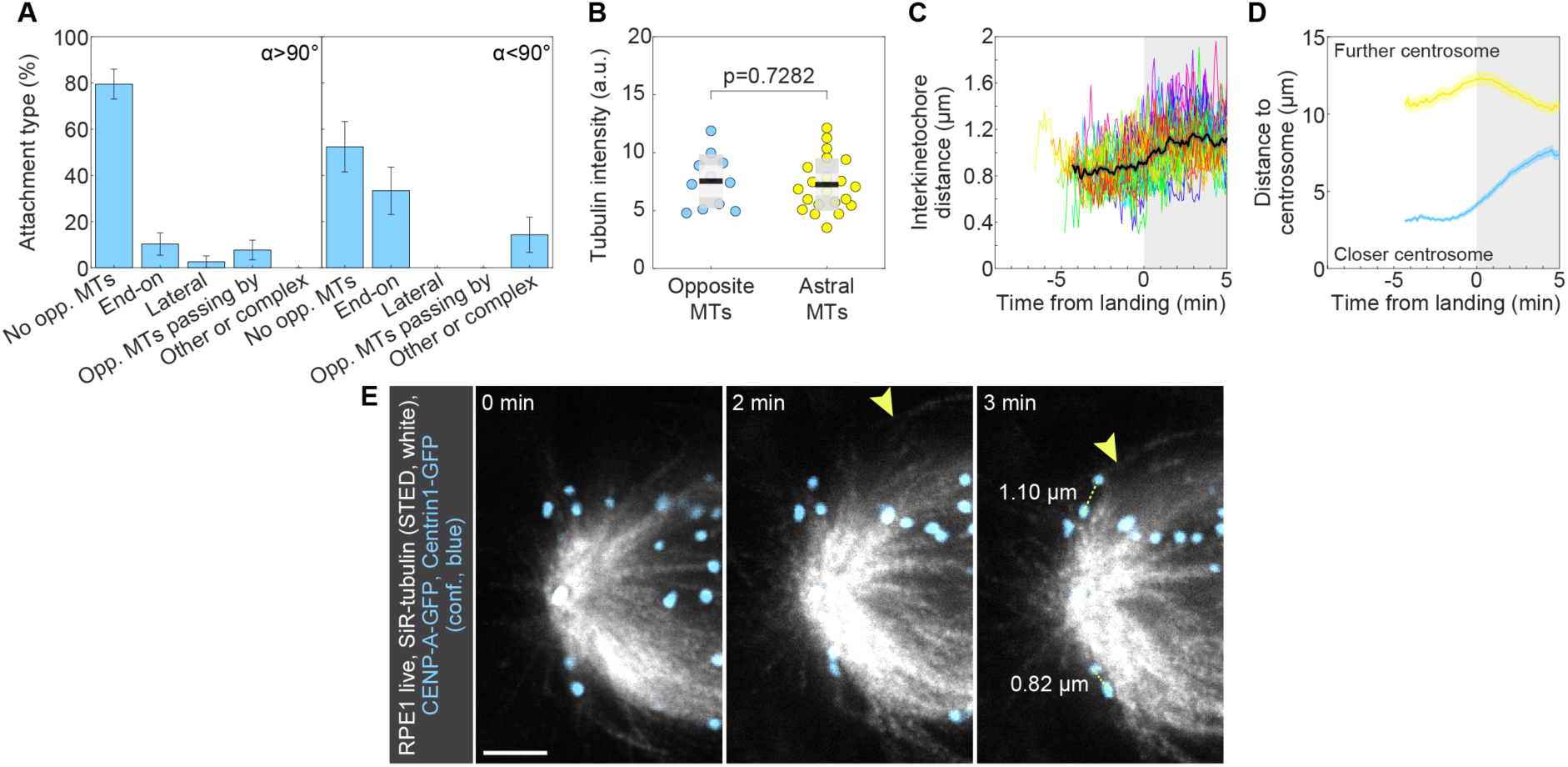
**(A)** Percentage of different attachment types with microtubules from the opposite side of the spindle for kinetochores between the end of centripetal movement and landing as measured from STED images in the areas with the pivoting angle above or below 90°. N = 8 kinetochore pairs in 8 cells from 3 experiments for above 90°. N = 10 kinetochore pairs in 9 cells from 4 experiments for below 90°. **(B)** Univariate scatter plot of the tubulin intensity for microtubules that come from the opposite side of the spindle and attach to polar kinetochores (blue) and astral microtubules (yellow). Boxes represent the standard deviation (dark gray), 95% confidence interval of the mean (light gray), and mean value (black). N = 11 opposite microtubules in 10 cells from 4 experiments. N = 22 astral microtubules in the same 10 cells from 4 experiments. Statistical test: t-test. **(C)** Interkinetochore distance of polar kinetochores over time from the end of centripetal movement. Different colors correspond to different kinetochore pairs. N = 39 kinetochore pairs in 20 cells from 8 experiments. **(D)** Distance to further (yellow) and closer (blue) centrosome over time for polar kinetochores from the end of centripetal movement. Values are shown as mean (dark line) and SEM (shaded areas). N = 39 kinetochore pairs in 20 cells from 8 experiments. **(E)** Time-lapse images from live-cell STED microscopy of RPE1 cells stably expressing CENP-A-GFP and Centrin1-GFP (blue, confocal) and stained with 100 nM SiR-tubulin (STED, white). Each image represents a single z-plane. The yellow arrow denotes the microtubule coming from the opposite side of the spindle. Dashed yellow lines represent the interkinetochore distance at landing. **(E)** Scale bar, 2 µm.

**Supplementary Figure 6.**
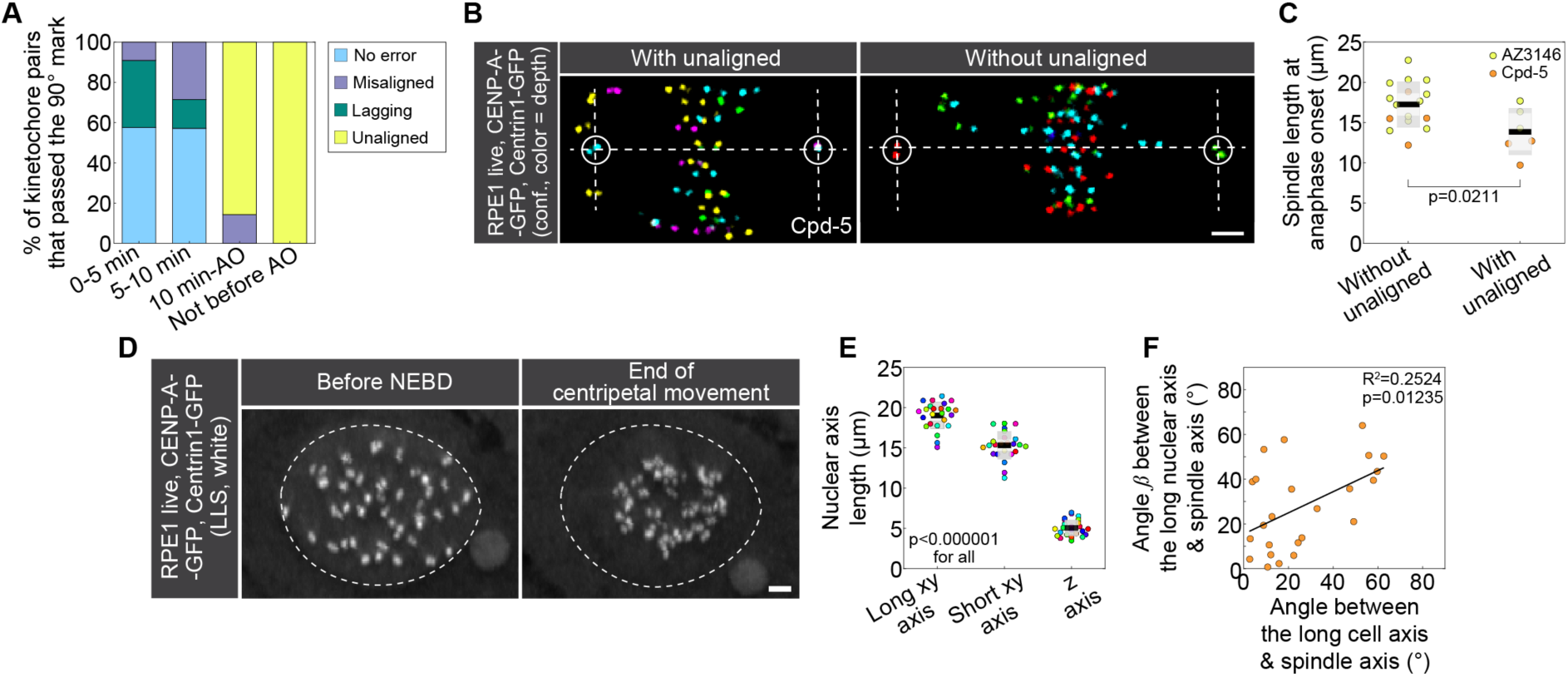
**(A)** Fractions of kinetochore pairs that passed the 90° mark in the first five minutes after NEBD, between 5 to 10 minutes after NEBD, between 10 minutes after NEBD and the anaphase onset, and those that failed to pass the 90° mark before anaphase onset. Kinetochore pairs are classified as no error (blue), lagging (green), misaligned (purple) and unaligned (yellow). N = 52 kinetochore pairs initially located behind the pole in 7 cells from 4 experiments. **(B)** Images of RPE1 cells stably expressing CENP-A-GFP and Centrin1-GFP (colorful, confocal) and treated with Mps1 inhibitor Cpd-5. Images show maximum intensity projections of the imaged cells. The left panel represents cells with unaligned chromosomes and the right panel represents cells without unaligned chromosomes at anaphase onset. Circles depict the spindle poles and dashed lines the boundaries of the polar region. Kinetochores are color-coded for depth with the Spectrum LUT in ImageJ throughout 4 z-planes, corresponding to 4 µm. Cells were originally imaged as part of the dataset for (Klaasen et al., 2022). **(C)** Univariate scatter plot of the spindle length at anaphase onset for cells without (left) and with (right) unaligned chromosomes at anaphase onset. Cells are divided based on the treatment, to those treated with AZ3146 (yellow) or Cpd-5 (orange). N = 7 cells from 4 experiments for AZ3146. N = 15 cells from 4 experiments for Cpd-5. Statistical test: t-test. Cells with added Cpd-5 were originally imaged as part of the dataset for (Klaasen et al., 2022). **(D)** Time-lapse images of RPE1 cells stably expressing CENP-A-GFP and Centrin1-GFP (white, lattice-lightsheet). Images show maximum intensity projections of 36 and 42 z-planes covering the whole cell, corresponding to 5.2 µm and 6.1 µm, respectively. The left image represents the nucleus before NEBD and the right image represents the kinetochore and centrosome arrangement at the end of centripetal movement. Dashed lines denote the borders of the nucleus as outlined before NEBD. **(E)** Univariate scatter plot of the long xy nuclear axis length (left), short xy nuclear axis length (middle) and z nuclear axis length (right). N = 24 cells from 8 experiments. Statistical test: Anova with post-hoc Tukey test. **(F)** A correlation between the angle that the long nuclear axis in interphase forms with the spindle axis at the end of the centripetal movement and the angle that the long cellular axis in interphase forms with the spindle axis at the end of the centripetal movement. N = 24 cells from 8 experiments. Linear regression. **(C,E)** Boxes represent standard deviation (dark gray), 95% confidence interval of the mean (light gray) and mean value (black). **(B,D)** Scale bar, 2 µm.

**Supplementary Figure 7.**
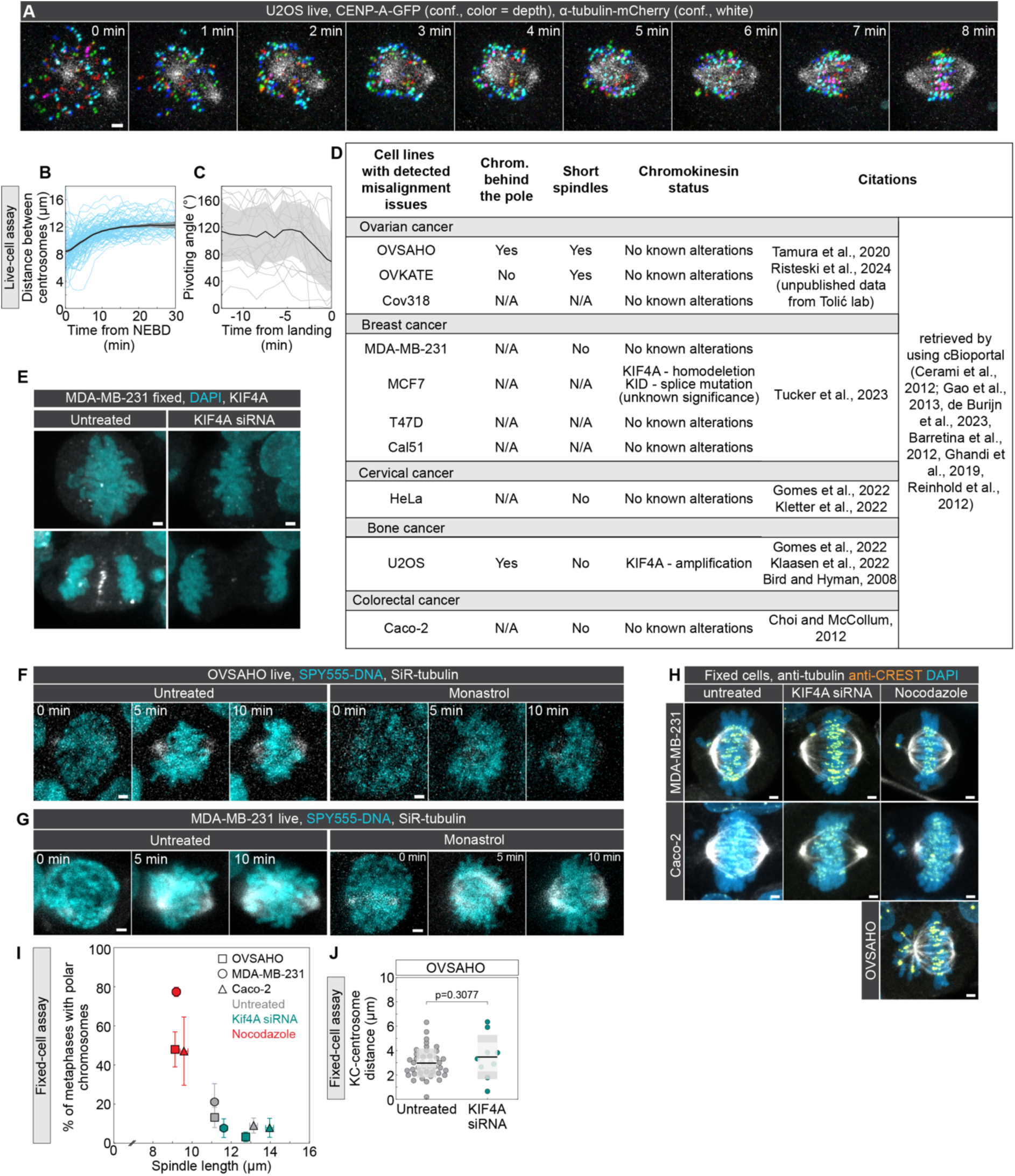
**(A)** Time-lapse confocal images of a live U2OS cell stably expressing CENP-A-GFP (colored for depth) and α-tubulin-mCherry (white), showing how polar kinetochores transit across the pole during spindle elongation. **(B)** Spindle length over time for U2OS cells as in **(A)**, N = 115 cells. Thick line, mean; shaded area, SEM; thin lines, individual cells. **(C)** Pivoting angle of pairs of unaligned kinetochores over time until landing, where time 0 denotes landing on the main spindle surface calculated for each kinetochore pair, for U2OS cells as in **(A)**, N = 23 kinetochore pairs from 23 cells. Thick line, mean; shaded area, SEM; thin lines, individual kinetochore pairs. **(D)** Mini-screen of cancer cell lines for studying elongation-driven pivoting, from the literature (Barretina et al., 2012; Bird and Hyman, 2008; Cerami et al., 2012; Choi and McCollum, 2012; de Bruijn et al., 2023; Gao et al., 2013; Ghandi et al., 2019; Gomes et al., 2022; Klaasen et al., 2022; Kletter et al., 2022; Reinhold et al., 2012; Risteski et al., 2024; Tamura et al., 2020; Tucker et al., 2023). **(E)** Images of fixed MDA-MB-231 cells, control (left) and treated with Kif4A siRNA (right), immunostained for Kif4A (white) and stained with DAPI (blue). **(F)** Time-lapse images of control (left) and treated with 15 µM monastrol (right) live OVSAHO cells stained with SPY-555-DNA (blue) and SiR-tubulin (white), imaged using lattice light-sheet microscopy. **(G)** Time-lapse images of control (left) and treated with 30 µM monastrol (right) live MDA-MB-231 cells stained with SPY-555-DNA (blue) and SiR-tubulin (white), imaged using lattice light-sheet microscopy. **(H)** Images of fixed immunolabeled cells of control (left column), KIF4A siRNA-treated (middle column), and nocodazole-treated (10 ng/mL, right column) MDA-MB-231 cells (top row), Caco-2 (middle row), and OVSAHO (bottom image). Cell are stained with anti-α-tubulin antibody (white), anti-CREST (orange) and DAPI (blue). **(I)** Fraction of fixed metaphase cells with polar chromosomes, i.e., at least one chromosome behind the pole, as a function of spindle length for fixed cells of 3 cancer cell lines (marked with different symbols as indicated). In the order of untreated (grey), Kif4A siRNA-treated (blue), and nocodazole-treated (10 ng/mL, red): N= 319, 157, 146 for OVSAHO, N= 128, 52, 115 for MDA-MB-231, and N= 122, 89, 138 for Caco-2. Error bars denote SEM. **(J)** Univariate scatter plot of the distance between the closer centrosome and the kinetochores of polar chromosomes for untreated (left) and KIF4A siRNA treated (right) fixed OVSAHO cells in metaphase. N = 39 chromosomes in 25 untreated cells from 3 experiments. N = 7 chromosomes in 6 KIF4A siRNA-treated cells from 2 experiments. Statistical test: t-test. Boxes represent standard deviation (dark gray), 95% confidence interval of the mean (light gray) and mean value (black). All scale bars, 2 µm.

### Supplementary Videos

**Video 1.** RPE1 cells stably expressing CENP-A-GFP and Centrin1-GFP (colorful, lattice light-sheet). Kinetochores are color-coded for depth with the brgbcmyw LUT in ImageJ throughout 32-40 z-planes. One polar kinetochore (white arrow) is tracked from NEBD until alignment to the spindle equator. Note that its passage across the polar region happens during the rapid movement of centrosomes (white circles) outwards. Scale bar, 2 µm.

**Video 2.** RPE1 cells stably expressing EYFP-tubulin (white, confocal) and H2B-mRFP (not imaged) with added HaloTag-CENP-A (blue, confocal). Polar kinetochore (white arrow) is adjacent to the microtubule signal that pivots around the spindle pole towards the spindle surface. Scale bar, 2 µm.

**Video 3.** RPE1 cells stably expressing EYFP-tubulin (white, confocal) and H2B-mRFP (not imaged) with added HaloTag-CENP-A (blue, confocal). Polar kinetochore (white arrow) is adjacent to the microtubule signal that pivots around the spindle pole towards the spindle surface. Scale bar, 2 µm.

**Video 4.** RPE1 cells stably expressing EB3-GFP (white) and H2B-mCherry (blue). Note that spindle poles are moving rapidly outwards, while chromosomes remain stationary during this process. Scale bar, 2 µm.

**Video 5.** RPE1 cells stably expressing CENP-A-GFP and Centrin1-GFP (colorful, lattice light-sheet) in DMSO control (left), STLC (middle) and FCPT (left) treatment. Kinetochores are color-coded for depth with the brgbcmyw LUT in ImageJ throughout 14-21 z-planes. In DMSO control cell, centrosomes (white circles) moved outwards, while polar kinetochore (white arrow) pivoted towards the spindle body. The situation was opposite in STLC treatment, where centrosomes moved inwards and polar kinetochore pivoted outwards. In FCPT treatment, spindle elongation was blockes and pivoting of polar kinetochore did not occur. Scale bar, 2 µm.

**Video 6.** RPE1 cells stably expressing CENP-A-GFP and Centrin1-GFP (colorful, lattice light-sheet) in monastrol treatment (left, duration 3 min 10 s) and after monastrol washout (right, duration 10 min 10 s). Kinetochores are color-coded for depth with the brgbcmyw LUT in ImageJ throughout 14-21 z-planes. Pivoting of polar kinetochore (white arrow) is driven by movement of centrosomes (white circle). Scale bar, 2 µm.

**Video 7.** RPE1 cells stably expressing CENP-A-GFP and Centrin1-GFP (colorful, lattice light-sheet) and treated with Mps1 inhibitor AZ3146. The left movie represents cells without unaligned chromosomes and the right movie represents cells with unaligned chromosomes. Kinetochores are color-coded for depth with the brgbcmyw LUT in ImageJ throughout 14-21 z-planes. White circles and arrows represent centrosomes and unaligned kinetochores, respectively. Scale bar, 2 µm.

**Video 8.** RPE1 cells stably expressing CENP-A-GFP and Centrin1-GFP (white, lattice light-sheet). Note that the cell aligned the centrosomes along the long nuclear and cellular axis. Scale bar, 2 µm.

**Video 9.** U2OS cell stably expressing EB3-GFP (white) and CENP-A-mCherry (blue) in early prometaphase, showing a region close to one centrosome (big white spot). Note that astral microtubules attached to a pair of polar kinetochores pivot around the centrosome towards the main spindle surface (in the clockwise direction). Time is given in min:s. Scale bar, 2 µm.

**Video 10.** U2OS cell stably expressing CENP-A–GFP (color-coded by depth, left panel) and α-tubulin–mCherry (white, right panel), imaged from nuclear envelope breakdown until telophase. Note that all polar kinetochores pivot around and successfully pass the centrosome early in mitosis. Kinetochores are depth color-coded using the 16-color LUT across all acquired z-planes. Scale bar, 2 μm.

**Video 11.** MDA-MB-231 cells stained with SPY-555-DNA (blue) and SiR-tubulin (white) imaged using lattice light-sheet microscopy. The left movie shows untreated cells, and the right movie shows cells treated with 30 µM monastrol. Note that the monastrol-treated cell has a delayed and reduced spindle elongation during prometaphase, leading to a delay in the movement of polar chromosomes. Scale bar, 2 µm.

**Video 12.** OVSAHO cells stained with SPY-555-DNA (blue) and SiR-tubulin (white) imaged using lattice light-sheet microscopy. The left movie shows an untreated cell with chromosomes stuck at the spindle pole in metaphase (white arrow), and the right movie shows a Kif4A siRNA-treated cell. Note that in the Kif4A siRNA-treated cell the spindle elongated more rapidly, rescuing the phenotype observed in the untreated cell. Scale bar, 2 µm.

## Notes

### Competing Interest Statement

The authors have declared no competing interest.

### Summary of Updates

1) To generalize our conclusions in other cell lines, we performed new experiments on four cancer cell lines: U20S (osteosarcoma), OVSAHO (ovarian carcinoma), MDA-MB-231 (triple-negative breast cancer), and Caco-2 (colorectal adenocarcinoma). Our new results show that microtubule pivoting occurs during spindle elongation in cancer cells, as in non-transformed cells, and facilitates successful congression of most polar kinetochores. In cancer cells, failure of timely polar chromosome transit during spindle elongation prolongs mitosis. When we slowed down spindle elongation by a low dose of monastrol, the resolution of polar chromosomes was delayed, whereas when we enhanced elongation by Kif4A depletion, polar chromosomes resolution was accelerated, in agreement with our model. These new results, presented in Figure 8 and Supplementary Figure 7, demonstrate that the spindle elongation-driven pivoting model applies across multiple cancer cell lines and plays a key role in ensuring rapid mitosis. 2) To characterize the attachment status of polar kinetochore-microtubule attachments, we tested Astrin as a positive marker of end-on attachments, together with Mad2. Moreover, we examined CENP-E and ZW10, components of the fibrous corona that promotes lateral attachments. Our new results show that kinetochores on polar chromosomes recruit fibrous corona proteins and Mad2, but lack Astrin. Non-polar kinetochores on prometaphase spindles recruit less corona proteins and less Mad2 than polar kinetochores, and still lack Astrin. This is different than in metaphase, where kinetochores recruit high levels of Astrin, low levels of corona proteins and lack Mad2. Together with our STED images, these new results imply that in early intact mitosis, both sister kinetochores on polar chromosomes typically form lateral attachments, while the sister closer to the pole additionally establishes immature or transient end-on attachments, which mature and stabilize as the kinetochores interact with the spindle body and congress. We added a new figure (Figure 5) to present these new results.

